# Systematic evaluation of structural connectome thresholding in whole-brain network modelling

**DOI:** 10.64898/2026.09.01.748520

**Authors:** Liyi Qian, Sihan Ju, Zhenyu Li, Yongsheng Xie, Qi Yue, Liang Chen

**Affiliations:** Department of Neurosurgery, Huashan Hospital, Fudan University, National Center for Neurological Disorders, Shanghai, China; Centre for Population Neuroscience and Stratified Medicine (PONS), Institute of Science and Technology for Brain-Inspired Intelligence, Fudan University, Shanghai, China

**Keywords:** structural connectome, thresholding, whole-brain network models, functional connectivity, functional connectivity dynamics

## Abstract

Large-scale whole-brain network models rely on structural connectomes to constrain simulated neural dynamics, yet the optimal processing of these anatomical scaffolds is not fully established. Here, we investigate the impact of structural connectome thresholding on whole-brain model performance to advance precision individual modelling. We measure model goodness-of-fit to both static functional connectivity and dynamic functional connectivity across a comprehensive spectrum of network densities, spatial parcellations, and model complexities using neuroimaging datasets of healthy individuals and a clinical cohort of post-stroke patients. We demonstrate that the optimal structural sparsity is highly resolution dependent. In coarse-grained parcellations, proportional thresholding enhances the model’s fit to static functional connectivity but relies on denser connectome to maintain dynamic functional fits. Conversely, fine-grained models require stringent connectome thresholding which simultaneously optimizes both static and dynamic functional fits. Taken together, these results indicate that the uncritical use of raw structural connectomes introduces suboptimal dynamical regimes, establishing resolution-tailored thresholding as an indispensable step for constructing precision brain network models.

## 1 Introduction

Large-scale whole-brain network models (BNMs) serve as a powerful framework to bridge the gap between brain structure and function (Breakspear, 2017; Deco et al., 2008; Deco & Kringelbach, 2014). While purely data-driven approaches exist (Luo et al., 2025), most biophysically grounded BNMs inherently rely on the structural connectome (SC) as an anatomical scaffold to simulate the functional patterns of brain using a minimal set of biophysical parameters (Deco & Kringelbach, 2014; H. E. Wang et al., 2024). BNMs not only simulate static functional connectivity (FC) as a manipulable and comparative biomarker, but also capture time-varying functional connectivity dynamics (FCD), providing temporal insights inaccessible through static methods (E. C. A. Hansen et al., 2015) to characterize brain state transitions across diverse physiological and pathological conditions (Deco et al., 2018, 2026; E. C. A. Hansen et al., 2015; V. K. Jirsa et al., 2014; Luppi et al., 2022). Recently, the field has increasingly shifted from investigating group-level universal mechanisms toward highly individualized modelling. This paradigm shift enables researchers to infer personalized dynamic properties, design precision neuromodulation interventions and predict clinical outcomes (An et al., 2022; V. Jirsa et al., 2023; Makhalova et al., 2022; Proix et al., 2017, 2018; Saberi, Wischnewski, et al., 2025; Zhang et al., 2026). However, the transition to single-subject inference imposes stringent demands on the reliability of the underlying biological constraints.

For BNMs, the fundamental biological constraint is provided by the SC, typically inferred from diffusion magnetic resonance imaging (dMRI). While group-level modelling can rely on SCs derived from large-scale, high-quality cohorts (Kong et al., 2021; Pfeffer et al., 2021; P. Wang et al., 2019) and further benefit from rigorously validated consensus construction frameworks (Betzel et al., 2019; Chang et al., 2023), individual-level SCs reconstructed from scans with variable imaging quality and acquisition protocols exhibit heightened susceptibility to the inherent limitations of dMRI, such as the ill-posed problem of disentangling intersecting fibre bundles (Maier-Hein et al., 2017; Sotiropoulos & Zalesky, 2019; C.-H. Yeh et al., 2021). Although state-of-the-art tractography algorithms have significantly enhanced sensitivity in resolving complex fibre configurations and reduced false-negative omissions (Farquharson et al., 2013; Tahedl et al., 2025; J. Tournier et al., 2012), this improvement comes at the cost of reduced specificity (Sarwar et al., 2019), often yielding nearly fully connected matrices. Previous studies indicate that when imaging quality is limited—a common scenario in clinical practice—this lack of specificity becomes even more pronounced (Seider et al., 2022).

Given the theoretical constraints of wiring costs and the necessity for efficient communication (Bullmore & Sporns, 2009; Rubinov & Sporns, 2010) as well as anatomical evidence from primates (Young, 1992), true brain connectivity is considered to be sparse. Although weak connections in tractography outputs do encompass some genuine long-range fibres (Ercsey-Ravasz et al., 2013), they predominantly consist of spurious connections (Thomas et al., 2014). Therefore, a straightforward and practical approach is to apply absolute or proportional thresholding to prune most weak edges (Fornito et al., 2016), allowing the SC to more accurately reflect the underlying biological network. In graph theoretical analysis, another SC-centric methodology, the impacts of such simple thresholding pre-processing have been extensively explored (Buchanan et al., 2020; Dimitriadis et al., 2021; Messaritaki et al., 2019). However, it remains largely unexplored in large scale whole-brain dynamic modelling. Current studies often adopt highly conservative approaches, either directly inputting the raw, nearly fully connected SC matrices (Demirtaş et al., 2019; Saberi, Wischnewski, et al., 2025) or applying arbitrary, minimal streamline cutoffs that barely reduce the overall network density (Deco et al., 2021; Rocha et al., 2022). This conservatism may stem partly from the fact that an overly sparse SC prevents some models from reaching a phase transition (Zarepour et al., 2019), and long-range connections play a crucial role in shaping human brain dynamics (Vohryzek et al., 2025). In spite of the recognized need to evaluate how thresholding affects model fits (Deco et al., 2021), conducting a detailed benchmark to evaluate the impact of sparsity on simulation outcomes and to identify the optimal strategy has previously been unfeasible due to the exorbitant computational costs associated with simulating non-linear dynamics across comprehensive parameter spaces. Consequently, it remains uncertain whether constructing whole-brain biophysical models based on near-dense SC matrices introduces systematic biases.

To address this critical gap, we systematically investigate the impact of structural connectome thresholding on the performance of individualized whole- brain network models. We first benchmarked the efficacy of proportional, absolute, and Orthogonal Minimum Spanning Tree (OMST) thresholding strategies. Specifically, we evaluated their impact on model goodness-of-fit across both static functional connectivity and time-varying functional connectivity dynamics. We subsequently validated the persistence of these effects in a clinical cohort of patients with post-stroke motor impairment. Furthermore, we extended our evaluation across diverse model architectures, ranging from the phenomenological Kuramoto model to Dynamic Mean-Field (DMF) models with increasing biophysical complexity and regional heterogeneity, while also tracking how structural sparsity inherently constrains the inferred biophysical parameter space. Finally, we examined the interplay between network resolution and structural sparsity across a wide range of spatial parcellation scales and explored the scale-dependent associations between structural graph-theoretical topology and simulated whole-brain dynamics.

## 2 Methods

### 2.1 Participants and data acquisition

#### 2.1.1 MICA-MICs dataset

The primary evaluation was conducted using the MICA-MICs dataset (Royer et al., 2022), comprising multimodal MRI data from healthy adults (*N*=50, 29.54 ± 5.62 years).

Imaging data were acquired using a 3T Siemens Magnetom Prisma-Fit scanner equipped with a 64-channel head coil. High-resolution structural T1-weighted images were obtained using a 3D MP-RAGE sequence (TR = 2300 ms, TE = 3.14 ms, TI = 900 ms, flip angle = 9°, voxel size = 0.8 mm isotropic). Resting-state fMRI (rs-fMRI) data were collected using a multiband accelerated 2D- BOLD echo-planar imaging (EPI) sequence (TR = 600 ms, TE = 30 ms, flip angle = 52°, voxel size = 3 mm isotropic, multiband factor = 6), during which participants were instructed to maintain fixation on a crosshair. Diffusion-weighted MRI (dMRI) was performed using a multi-shell EPI sequence (TR = 3500 ms, TE = 64.4 ms, voxel size = 1.6 mm isotropic, multiband factor = 3). The diffusion protocol consisted of three shells with b-values of 300, 700, and 200 s/mm², with 10, 40, and 90 directions per shell, respectively. Specific details regarding the acquisition protocols refer to (Royer et al., 2022).

#### 2.1.2 HCP-YA dataset

To evaluate modelling based on group-level SC, we used a consensus structural connectome derived from 706 healthy participants from the Human Connectome Project Young Adult (HCP-YA) cohort. Detailed descriptions can be found in previous publications (Glasser et al., 2013, 2016; Van Essen et al., 2013).

#### 2.1.3 Post-stroke motor impairment dataset

We validated our findings in a cohort of 15 stroke patients. The mean age of the patients was 54.27 years (SD = 11.96). All patients were formally diagnosed with ischemic stroke. All participants were right-handed and presented with unilateral hemiplegia. Data acquisition, analysis, and publication of the results were approved by the Ethics Committee of Huashan Hospital, Fudan University (No. KY2022-676). Written informed consent was obtained from all patients prior to their participation in the study.

All neuroimaging data were collected at Huashan Hospital, Fudan University, utilizing a 3.0 Tesla UIH uMR 790 scanner (United Imaging Healthcare, Shanghai, China). High-resolution 3D T1-weighted structural images were acquired using a gradient-echo sequence. Scanning parameters included a TR of 7.2 ms, a TE of 2.3 ms, a TI of 1000 ms, an 8° flip angle, and an isotropic slice thickness of 1.0 mm. Resting-state BOLD data were obtained using a 2D EPI sequence. To achieve high temporal resolution, a multiband acceleration factor of 7 was applied. The acquisition parameters consisted of a TR of 700 ms, a TE of 33 ms, a 52° flip angle, and a slice thickness of 2.5 mm. A 2D EPI protocol featuring a multiband acceleration factor of 3 was employed for diffusion imaging. The scanning settings included a TR of 5300 ms, a TE of 82 ms, a 90° flip angle, and a 1.5-mm slice thickness. A multi-shell paradigm was implemented, incorporating b-values of 0, 1000, and 2000 s/mm². Specifically, four unweighted volumes (b = 0 s/mm²) were collected to facilitate motion correction and establish a signal baseline, while gradients for the active diffusion weightings were distributed across 32 distinct spatial directions.

### 2.2 Image preprocessing and connectome construction

#### 2.2.1 Diffusion weighted image processing

For the MICA-MICs dataset, structural connectomes were generated using the micapipe pipeline (Cruces et al., 2022), which employs MRtrix3 (J.-D. Tournier et al., 2019) to perform multi-shell multi-tissue constrained spherical deconvolution and SIFT2 (Smith et al., 2015)-weighted tractography (40 million streamlines). For comprehensive details please refer to the original publication (Royer et al., 2022).

Group-level SC was adopted from a previous study (Saberi, Wan, et al., 2025), derived from the HCP-YA dataset using an in-house pipeline (Jung et al., 2021).

For the post-stroke motor impairment dataset, dMRI was preprocessed via QSIPrep (v1.0.1) and reconstructed using the *mrtrix_multishell_msmt_ACT- hsvs* workflow in QSIRecon (v1.0.1) (Cieslak et al., 2021). Processing included MP-PCA denoising (Veraart et al., 2016), eddy current correction (Andersson & Sotiropoulos, 2016), and MSMT-CSD (Jeurissen et al., 2014) to estimate fibre orientation distributions. Whole-brain probabilistic tractography was performed using the iFOD2 algorithm (Calamante & Connelly, 2010) to generate 10^7^ streamlines with anatomical constraints enforced by HSVS. These streamlines were subsequently weighted by SIFT2 (Smith et al., 2015). A detailed, reproducible description of the pipelines and all associated software citations are provided in Supplementary Methods S1 and S2.

#### 2.2.2 Functional imaging processing

For the MICA-MICs dataset, rs-fMRI data were processed using micapipe (Cruces et al., 2022), including motion and distortion correction. Nuisance signals were removed via ICA-FIX (Salimi-Khorshidi et al., 2014) and spike regression. Denoised time series were co-registered to T1w space, mapped to the individual’s cortical surface, and underwent spatial smoothing with a 10 mm FWHM Gaussian kernel.

For the post-stroke motor impairment dataset, T1w and rs-fMRI data were preprocessed using fMRIPrep (v24.1.1) (Esteban et al., 2019). A detailed, reproducible description of the pipelines and all associated software citations are provided in Supplementary Methods S3. For patients with visible stroke, manual masks delineated in ITK-SNAP (v4.4.0) (Yushkevich et al., 2006) were incorporated via cost-function masking during nonlinear registration to prevent spurious warping. The BOLD time-series were subjected to motion and susceptibility distortion correction, then normalized to MNI space. To mitigate lesion-driven artifacts and ensure accurate functional connectivity, an ICA-based denoising approach (Lamouroux et al., 2025) was subsequently applied to the preprocessed data. Further nuisance regression and temporal filtering (0.01–0.1 Hz) were implemented in Nilearn (v0.10.2). To prevent lesion-driven artifacts from contaminating adjacent healthy tissue, no spatial smoothing was applied to the functional data.

#### 2.2.3 Structural connectivity parcellation and thresholding

For each dataset, SC matrices were constructed by parcellating the whole-brain tractograms into multiple spatial resolutions. Cortical nodes were defined using the Desikan-Killiany (DK) atlas (68 nodes) (Desikan et al., 2006) and the Schaefer atlas at resolutions ranging from 100 to 300 parcels (Schaefer et al., 2018). Additionally, 14 subcortical nodes (Fischl et al., 2002) were extracted combined with Schaefer 100 atlas.

Proportional thresholding (Prop) was applied to maintain a fixed percentage of the strongest connections, ranging from 5% to 100% (no thresholding) with a step size of 5%. For higher resolutions (Schaefer 200–300), a finer Prop range of 0% to 10% was additionally evaluated with a 1% step size. Absolute thresholding (Abs) was performed by removing edges with streamline counts below 3, 5, 10, or 20. To derive a topology-driven backbone without an arbitrary weight cutoff, we utilized Orthogonal Minimum Spanning Trees (OMST) (Dimitriadis et al., 2017). This method iteratively extracts nested MSTs that maximize the information captured while minimizing redundant edges, providing a parameter-free approach to balance network sparseness and connectivity. Following previous studies (Saberi, Wan, et al., 2025, 2025), the SC matrices were normalized by dividing its mean × 100, resulting in an equal mean of 0.01 in all SCs.

#### 2.2.4 Empirical functional connectivity and dynamics construction

Static functional connectivity (FC) was computed as the Pearson correlation between the BOLD time series of all regional pairs. For functional connectivity dynamics (FCD), a sliding window approach was employed using a window length of 30 TR and a sliding step of 5 TR. A separate FC matrix was calculated for each temporal window, and the FCD matrix was subsequently generated by computing the Pearson correlation between the lower triangular elements of these windowed FC matrices.

### 2.3 Brain network modelling

All brain network simulations and parameter optimizations were implemented using cuBNM (v0.1.0) (Saberi, Wan, et al., 2025), a recently developed CUDA- accelerated framework designed for high-throughput brain network modelling.

#### 2.3.1 Dynamic Mean-Field model

The macroscopic neural dynamics of each parcellated brain region are simulated using the Dynamic Mean-Field (DMF) model (Deco et al., 2013; Deco, Ponce-Alvarez, et al., 2014).

In this framework, each network node *i* represents a local circuit consisting of coupled excitatory (*E*) and inhibitory (*I*) neuronal pools. The temporal evolution of this system is governed by a set of coupled stochastic differential equations. The total input currents to the excitatory and inhibitory populations, *I_E_*_,*i*_ and *I_I_*_,*i*_, are determined by the local recurrent connections and the global long-range connections scaled by the structural connectivity matrix *C_ij_*:

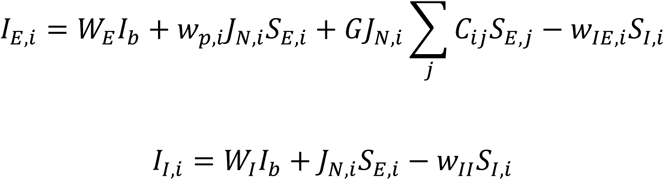

where *W_E_ I_b_* and *W_I_ I_b_* represent the background currents, *G* is the global coupling strength, *w_p_*_,*i*_ is the local recurrent excitatory connection weight, and *J_N_*_,*i*_ represents the NMDA receptor conductance. The resulting population firing rates, *r_E_*_,*i*_ and *r_I_*_,*i*_, are driven by these input currents through a non-linear input-output transfer function *H*:

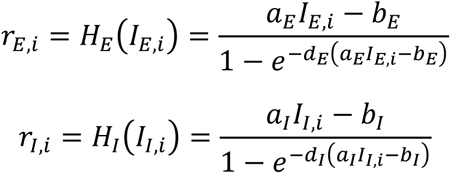

with *a_E_*_,*I*_, *b_E_*_,*I*_, and *d_E_*_,*I*_ serving as fixed physiological constants for the respective populations. The dynamics of the corresponding synaptic gating variables, *S_E_*_,*i*_ and *S_I_*_,*i*_, which mediate the interactions between populations across the network, evolve according to:

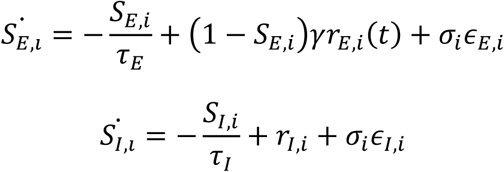

where *τ_E_* and *τ_I_* denote the synaptic time constants, *γ* is a kinetic parameter, and *ε* introduces uncorrelated standard Gaussian noise scaled by the noise amplitude *σ_i_* . To maintain biological plausibility, the inhibitory-to-excitatory connection weight (*w_IE_*_,*i*_) is determined via a Feedback Inhibition Control (FIC) algorithm (Deco, Ponce-Alvarez, et al., 2014), which regulates the average firing rate of the excitatory populations to a baseline of approximately 3 Hz.

The model setup consisted of three distinct parameterization schemes. In the first scheme, only the global coupling strength (*G*) was configured as a free variable, while the local recurrent excitatory connection weight (*w_p_*) and NMDA receptor conductance (*J_N_*) were set to 1.4 and 0.15 across all regions (Deco et al., 2018; Wong & Wang, 2006). In the second scheme, all three parameters (*G*, *w_p_*, and *J_N_*) were treated simultaneously as global free variables, maintaining the assumption of a homogeneous cortex (Saberi, Wan, et al., 2025). In the third scheme, spatial heterogeneity was introduced to the regional parameters *w_p_*_,*i*_ and *J_N_*_,*i*_ based on the T1w/T2w ratio (myelin map) (Demirtaş et al., 2019). For a given node *i*, the heterogeneous parameter *P_i_* (where *P* ∈ {*w_p_*, *J_N_*}) was defined as:

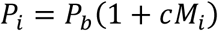

where *P_b_* represents the baseline value of the parameter, *c* is a scaling coefficient, and *M_i_* corresponds to the min-max normalized myelin map value for that specific region.

#### 2.3.2 Kuramoto model

In this phenomenological framework, the macroscopic dynamics of each parcellated brain region are represented by a phase oscillator, simulating the continuous phase synchronization dynamics of the network (Kuramoto, 1984). The temporal evolution of the phase θ*_i_* for each node *i* is governed by the stochastic differential equation:

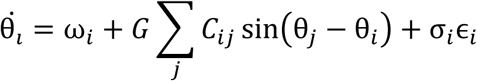

where ω*_i_* denotes the intrinsic natural frequency of the local oscillator, *C_ij_* represents the structural connectivity weight from node *j* to node *i*, and *G* serves as the global coupling strength that scales these inter-regional phase interactions. Additive standard Gaussian noise is introduced into the system via ɛ*_i_*, scaled by the local noise amplitude σ*_i_*.

#### 2.3.3 Balloon-Windkessel model

The simulated node-wise neural activity was transformed into observation-level Blood-Oxygen-Level-Dependent (BOLD) signals using the Balloon-Windkessel hemodynamic model (Buxton et al., 1998; Heinzle et al., 2016).

For each network node *i*, the simulated excitatory synaptic gating variable *S_E_*_,*i*_ served as the driving neural input to the regional hemodynamic response. The temporal evolution of this localized hemodynamic state is described by a system of differential equations governing the vasodilatory signal *x_i_*, blood inflow *f_i_*, blood volume *v_i_*, and deoxyhemoglobin content *q_i_*:

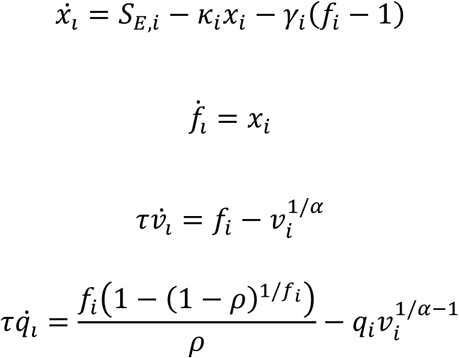

where *κ_i_* represents the rate of signal decay, *γ_i_* is the rate of flow-dependent elimination, *τ* denotes the hemodynamic transit time (set to 0.98 s), *α* is Grubb’s exponent (set to 0.32), and *ρ* represents the resting oxygen extraction fraction (set to 0.34). Following the numerical integration of these intermediate state variables, the final simulated BOLD signal for each node was computed at every prespecified TR step as:

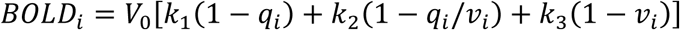

where *V*_0_ represents the resting blood volume fraction (0.02), and the coefficients *k*_1_, *k*_2_, and *k*_3_ are scanner-specific constants set to 3.72, 0.527, and 0.53, respectively, corresponding to a standard 3T magnetic resonance imaging acquisition protocol. The resulting simulated BOLD time series were subsequently utilized for the derivation and evaluation of functional connectivity metrics.

#### 2.3.4 Model performance metrics

The static functional connectivity fit ( *FC_corr_*) was defined as the Pearson correlation coefficient between the lower triangular elements of the simulated and empirical static functional connectivity matrices. To account for temporal transitions, functional connectivity dynamics (FCD) matrices were constructed by cross-correlating dynamic states across sliding time windows, utilizing parameters identical to those applied to the empirical data. The dynamic fit (*FCD_KS_*) was subsequently calculated as the Kolmogorov-Smirnov distance between the simulated and empirical FCD distributions (E. C. A. Hansen et al., 2015). Inter-hemispheric connections were excluded from all FC and FCD calculations to mitigate the influence of tracking inaccuracies associated with long-range callosal fibers (Messé et al., 2014), which can compromise the reliability of model performance.

While an exhaustive grid search treats the parameter space uniformly without an explicit optimization trajectory, the presence of high-dimensional parameters necessitates a cost-driven approach. Consequently, individual performance metrics were integrated into a unified objective function to be minimized during the optimization process, consistent with previous studies (Saberi, Wan, et al., 2025; Saberi, Wischnewski, et al., 2025):

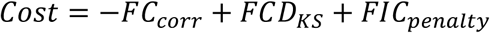

The *FIC_penalty_* term was incorporated to enforce biological realism through FIC.

#### 2.3.5 Model optimization

For the single-parameter configuration, a systematic grid search was performed across the global coupling strength (*G*). This search evaluated 300 equidistant points within the range of [0.001, 3].

For the three-parameter extension, the model incorporated the recurrent excitatory connection weight ( *w_p_*) and NMDA receptor conductance ( *J_N_*) alongside *G* . Optimization was conducted using the Covariance Matrix Adaptation Evolution Strategy (CMA-ES) (N. Hansen, 2016) with a population size of 128 and a maximum of 120 iterations.

For the incorporation of cortical heterogeneity, local biophysical variables were parameterized as functions of T1w/T2w-derived myelin maps from the HCP dataset. This spatial parameterization defined a joint optimization task for the global variables and their corresponding regional scaling coefficients, utilizing the CMA-ES algorithm to minimize the cost function until convergence.

### 2.4 Graph-theoretical metrics

To characterize the structural connectome, several graph-theoretical metrics were calculated based on the weighted adjacency matrix *W* and its corresponding cost matrix *L* = 1/*W*.

Modularity (*Q*) was estimated using the Louvain algorithm (Blondel et al., 2008) to quantify network segregation into non-overlapping communities. Global efficiency (*E_glob_*), an index of global integration, was defined as the average inverse shortest path length derived from the cost matrix *L* via Dijkstra’s algorithm. Local segregation was assessed using the weighted clustering coefficient (*C*) (Onnela et al., 2005), averaged across all nodes. Finally, small-world propensity (**φ**) (Muldoon et al., 2016) was computed to measure the deviation of observed clustering and path lengths from weighted lattice and random null models, with values closer to 1 indicating higher small-worldness. All metrics were implemented using the Brain Connectivity Toolbox (Rubinov & Sporns, 2010), netneurotools (v0.3.0) (Liu et al., 2025) and small-world- propensity (v0.0.19) (Muldoon et al., 2016).

### 2.5 Statistical analysis

All statistical analyses were conducted using SciPy (v1.14.1) and Statsmodels (v0.14.2). The normality of the distribution for all model performance metrics and topological indices was assessed using the Shapiro-Wilk test. Descriptive statistics for the model performance metrics and graph-theoretical measures were reported as mean ± standard error of the mean (SEM) across subjects. To evaluate the impact of varying SC thresholding strategies on model performance and topological metrics, paired Student’s t-tests were utilized to compare each sparsification level against the unthresholded SC. To explore the relationship between the network’s structural topology and the dynamic model’s fitting performance across different spatial scales, Spearman rank correlation coefficients were calculated between the SC graph-theoretical metrics and the DMF model performance metrics. Statistical significance was established at an alpha level of 0.05.

## 3 Results

### 3.1 Impacts of SC density on biophysical network model fitting

For each individual in the MICA-MICs cohort (*N*=50), subject-specific SC was extracted and parcellated using the Schaefer-100 atlas (Schaefer et al., 2018), which represents one of the most widely adopted spatial scales in whole-brain network modeling (Fig. 1A). While the raw SC reconstruction resulted in nearly fully dense networks, we systematically varied the network density from 0.05 to 1.0 to evaluate its impact on biophysical network modelling. Furthermore, we evaluated the efficacy of other common SC thresholding strategies. These included absolute streamline counts thresholds, which are frequently employed in whole-brain modelling (Rocha et al., 2022; Saberi, Wischnewski, et al., 2025), and the Orthogonal Minimum Spanning Tree (OMST) method (Dimitriadis et al., 2017), which is favoured in graph-theoretical studies for its ability to preserve core topological properties (Messaritaki et al., 2019). Representative SC matrices and their corresponding edge-length distributions across these thresholding strategies are presented in Supplementary Fig. 1. Note that absolute streamline count thresholding had only a minimal impact on the SC, whereas the OMST resulted in a density of approximately 0.03.

**Fig. 1.**
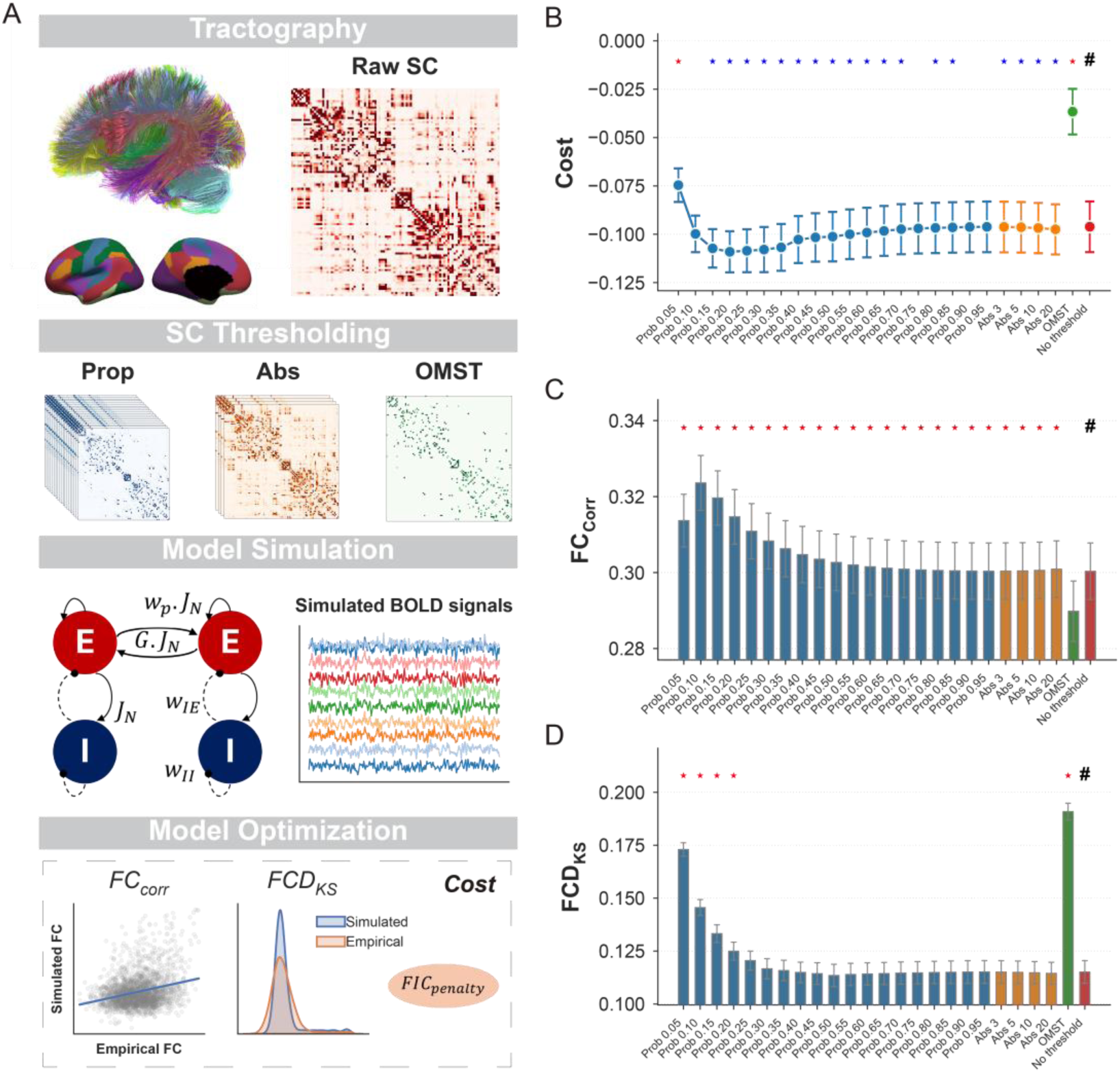
Impact of SC thresholding on biophysical network model performance. (A) Schematic of the modelling pipeline. Subject-specific raw SC matrices were processed using proportional (Prop), absolute (Abs), and Orthogonal Minimum Spanning Tree (OMST) thresholding strategies. The resulting networks served as structural backbones for Dynamic Mean Field model simulations. Model optimization was evaluated by comparing simulated and empirical BOLD signals to compute an integrated Cost metric, derived from correlation of static functional connectivity ( *FC_Corr_*), Kolmogorov-Smirnov distance of functional connectivity dynamics (*FCD_KS_*) and the feedback inhibition control penalty (*FIC_penalty_*). (B-D) Evaluation of model fitting across varying network densities and thresholding strategies, quantified by the integrated *Cost* metric (B), *FC_Corr_* (C), and *FCD_KS_* (D). The ’#’ symbol denotes the unthresholded baseline. Stars indicate statistically significant differences compared to the baseline, blue indicates a significant decrease, while red indicates a significant increase (*P* < 0.05, paired t-test). Data was presented as mean ± SEM.

Using these matrices of varying thresholding strategies as the structural backbone, we implemented the Dynamic Mean Field (DMF) model, where each node was represented by coupled excitatory and inhibitory neuronal pools (Deco et al., 2013; Deco, Ponce-Alvarez, et al., 2014). To identify the optimal dynamical regime and independently evaluate different optimization objectives, we performed a grid search across the global coupling parameter (*G*). Model performance was quantified using three primary metrics: an integrated *Cost* metric (Saberi, Wischnewski, et al., 2025), the Pearson correlation with static functional connectivity (*FC_Corr_*), and the Kolmogorov-Smirnov distance of the functional connectivity dynamics (*FCD_KS_*) (E. C. A. Hansen et al., 2015).

Fig. 1B illustrates that the integrated Cost metric exhibits an asymmetrical U-shaped trajectory. The metric decreases sharply from the lower density bound, reaches a global minimum at approximately 0.25, and subsequently increases as the networks approach full density. Notably, the absolute thresholding strategy had a negligible impact on the Cost metric. In contrast, the OMST approach led to a significant decline in model performance.

To elucidate the drivers underlying the observed U-shaped trend, we independently examined the effects of network density on the fitting accuracy of *FC_Corr_* and *FCD_KS_* . Interestingly, the two metrics exhibited markedly divergent density-dependent patterns (Fig. 1C-D). For *FC_Corr_*, we observed that even the most permissive absolute thresholding yielded improvements in fitting performance, with the accuracy peaking at a density of 0.1. Only the OMST failed to provide a significant enhancement over the unthresholded baseline. Conversely, *FCD_KS_* appeared more resilient to density variations, showing performance degradation only when the density dropped below 0.2 or when OMST was employed. Across other density levels, the *FCD_KS_* fitting performance remained statistically comparable to that of the unthresholded networks. Following the observation that proportional strategies outperformed both OMST and absolute thresholding, subsequent analyses were conducted solely using proportional thresholding.

To determine whether these observations were driven by properties of individual-level SC data or the dataset itself, we employed a common structural backbone from the HCP-YA dataset across all subjects. The modelling results remained highly consistent with the findings based on individual-level SC models, with only minor shifts in peak positions (Supplementary Fig. 2). Cost reached its minimum at approximately 0.2, and the fit for *FC_Corr_* was optimized at a relatively sparse density (0.15), while *FCD_KS_* showed comparable performance to unthresholded results at densities above 0.10.

### 3.2. Influence of SC density persist across pathological brain architectures

To assess whether these patterns persist in more complex clinical scenarios, we applied the same framework to a clinical cohort of patients with post-stroke motor impairment (*N*=15). This dataset involved a distinct imaging acquisition and preprocessing protocol, and the subjects presented with brain lesions (Fig. 2A-B).

**Fig. 2.**
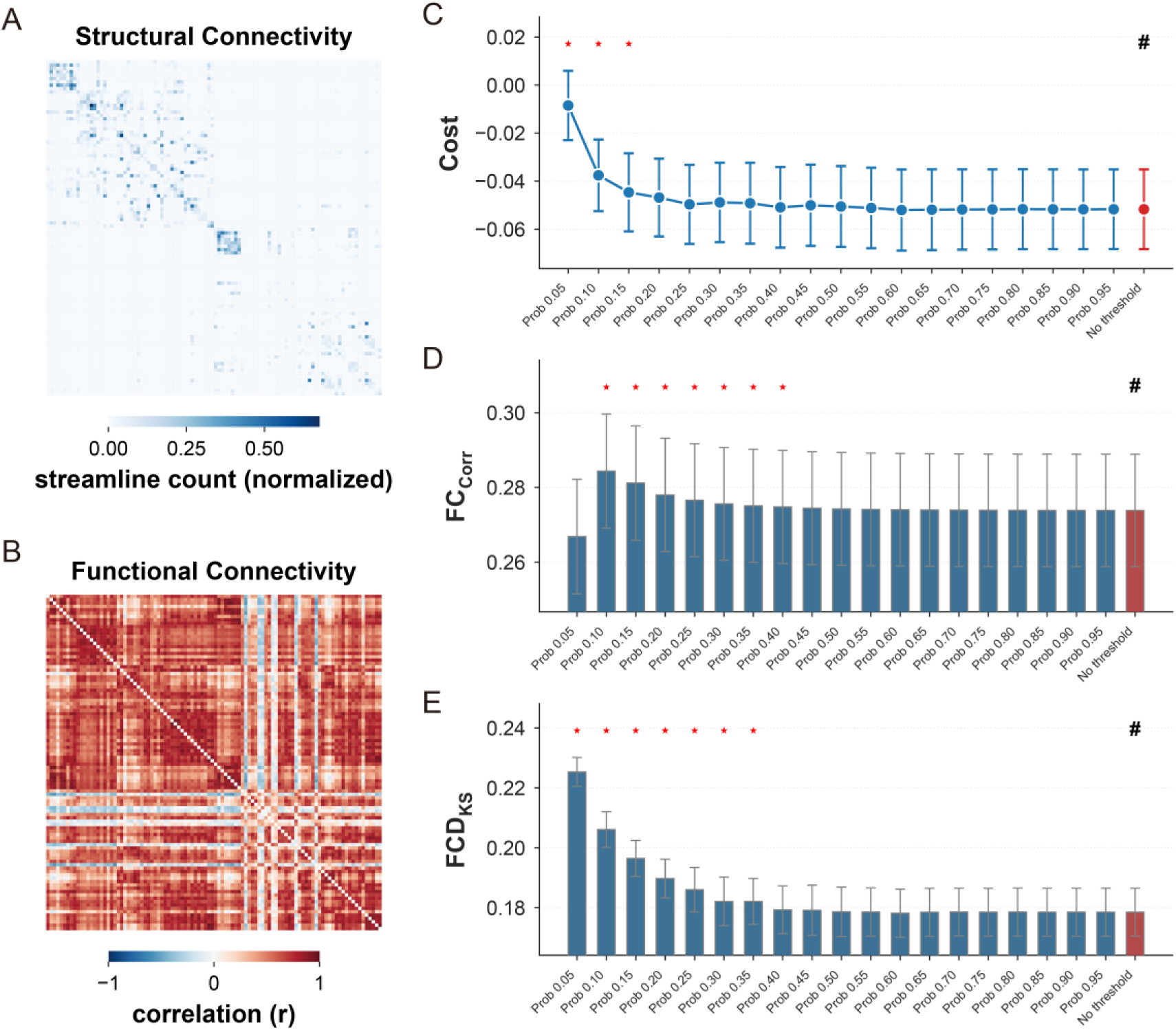
Influence of SC density on biophysical network model performance in stoke dataset. (A-B) Structural Connectivity (A) and Functional Connectivity (B) examples for the clinical cohort of patients with post-stroke motor impairment. (C-E) Evaluation of model fitting across varying network densities, quantified by the integrated Cost metric (C), *FC_Corr_* (D), and *FCD_KS_* (E). The ’#’ symbol denotes the unthresholded baseline. Stars indicate statistically significant differences compared to the baseline, blue indicates a significant decrease, while red indicates a significant increase (*P* < 0.05, paired t-test). Data was presented as mean ± SEM.

Although the integrated Cost metric did not exhibit a statistically significant minimum at low densities (Fig. 2C), independent evaluations of the static and dynamic functional fits revealed trajectories consistent with the healthy cohort (Fig. 2D-E). Specifically, the fit for *FC_Corr_* reached a distinct peak at a relatively sparse density (0.10) before gradually declining, whereas *FCD_KS_* performed poorly at high sparsity but reached a plateau once densities exceeded 0.40.

### 3.3. Impacts of SC density across diverse whole-brain modelling configurations

One recent trend in whole-brain modeling is the inclusion of additional degrees of freedom to enhance model performance. Having established the impact of SC density on the simplest model configuration, we next sought to determine whether these patterns hold when the model’s complexity is increased.

We first extended the DMF model by incorporating additional biophysical free parameters. Beyond the global coupling strength (*G*), we allowed the recurrent excitatory connection weight (*w_p_*) and NMDA receptor conductance (*J_N_*) to vary. Despite the increased degrees of freedom, which generally improved overall model performance, the underlying trends regarding SC density remained consistent (Fig. 3A-C). Specifically, *FC_Corr_* benefited from higher sparsity, whereas *FCD_KS_* required moderate densities to achieve performance comparable to unthresholded results.

**Fig. 3.**
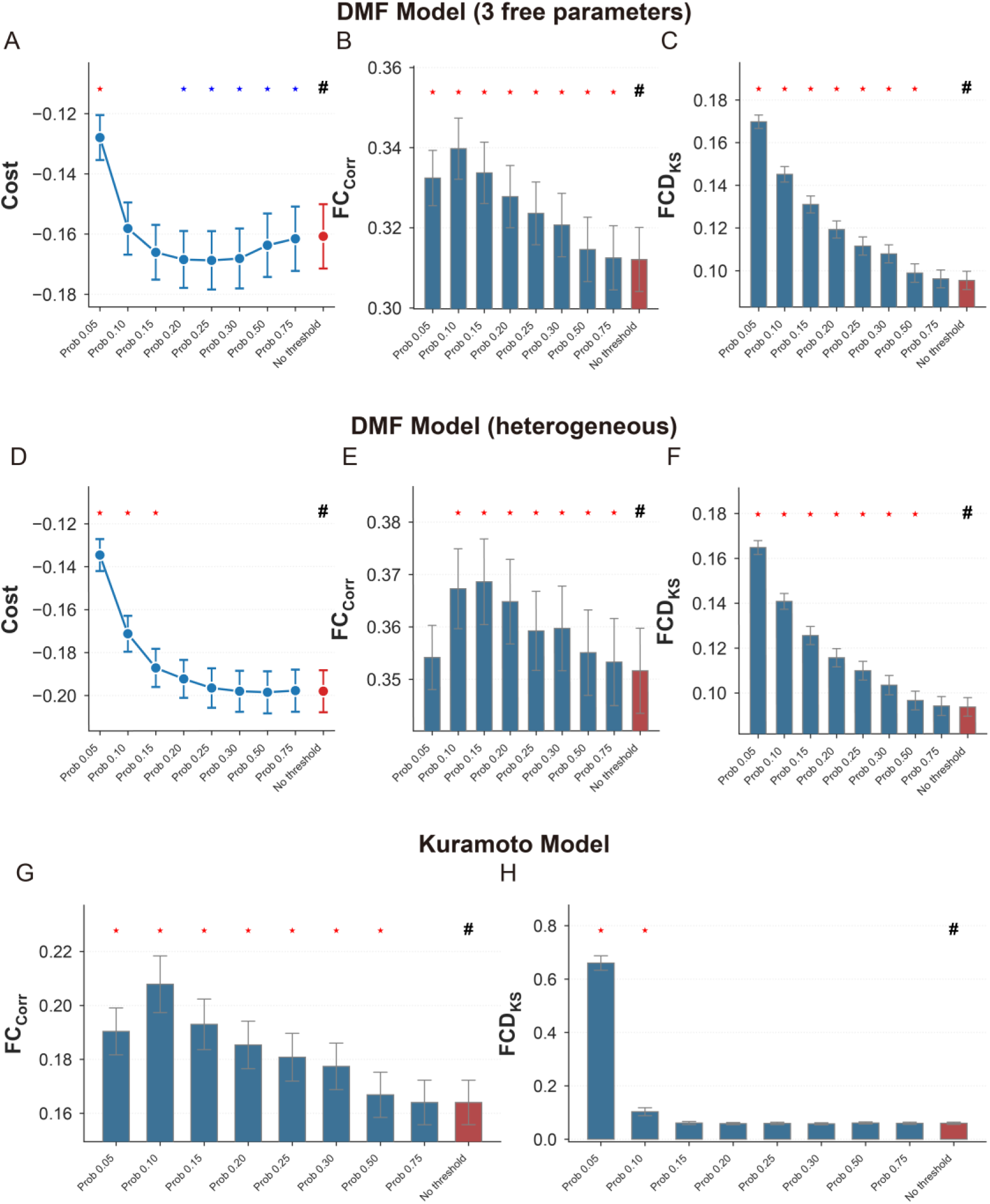
Impacts of SC density across diverse whole-brain modelling configurations. (A-C) Evaluation of model fitting for the extended DMF model with three free parameters (global coupling *G*, recurrent excitatory connection weight *w_p_*, and NMDA receptor conductance *J_N_*), quantified by the integrated *Cost* metric (A), *FC_Corr_* (B), and *FCD_KS_* (C) across varying network densities. (D-F) Evaluation of model fitting for the heterogeneous DMF model, where local biophysical parameters were scaled according to the T1w/T2w myelin map, quantified by Cost (D), *FC_Corr_* (E), and *FCD_KS_* (F). (G-H) Evaluation of model fitting for the Kuramoto oscillator model, quantified by *FC_Corr_* (G) and *FCD_KS_* (H). Stars indicate statistically significant differences compared to the baseline, blue indicates a significant decrease, while red indicates a significant increase (*P* < 0.05, paired t-test). Data was presented as mean ± SEM.

To further assess the robustness of our findings, we introduced regional heterogeneity by scaling the local biophysical parameters ( *w_p_* and *J_N_*) according to the T1w/T2w myelin map. This approach provides a biologically informed spatial prior that reflects the brain’s primary sensorimotor-to-association hierarchical gradient, thereby enhancing model performance with a minimal increase in free parameters. While this spatial heterogeneity further enhanced the overall fit to empirical data, the characteristic patterns of SC density’s impact on *FC_Corr_* and *FCD_KS_* remained similar (Fig. 3D-F).

Finally, to test whether these findings were tied to the specific mathematical framework of the DMF model, we replicated the evaluation using the Kuramoto oscillator model (Kuramoto, 1984), which simulates the phase synchronization dynamics of coupled oscillators. Sparse SC continues to facilitate better *FC* fitting in the Kuramoto model, with negative impacts on *FCD_KS_* appearing only when density dropped below 0.15 (Fig. 3G-H).

### 3.4. SC density as a constraint on the biophysical parameter space

Our results reveal that the impact of SC density on model performance is not overshadowed by improvements gained from additional degrees of freedom. Conversely, we sought to test whether the optimal parameters inferred at different densities exhibit systematic shifts, which can be interpret as a quantitative characterization of individual functional brain architecture.

In the single-parameter DMF model, the global coupling strength ( *G*) that yielded the best fit exhibited a monotonic increase as a function of network density (Fig. 4A). In the extended three-parameter model, the optimal *G* followed a similar upward trajectory with increasing density, whereas the recurrent excitatory connection weight (*w_p_*) and NMDA receptor conductance (*J_N_*) declined as density increased (Fig. 4B).

**Fig. 4.**
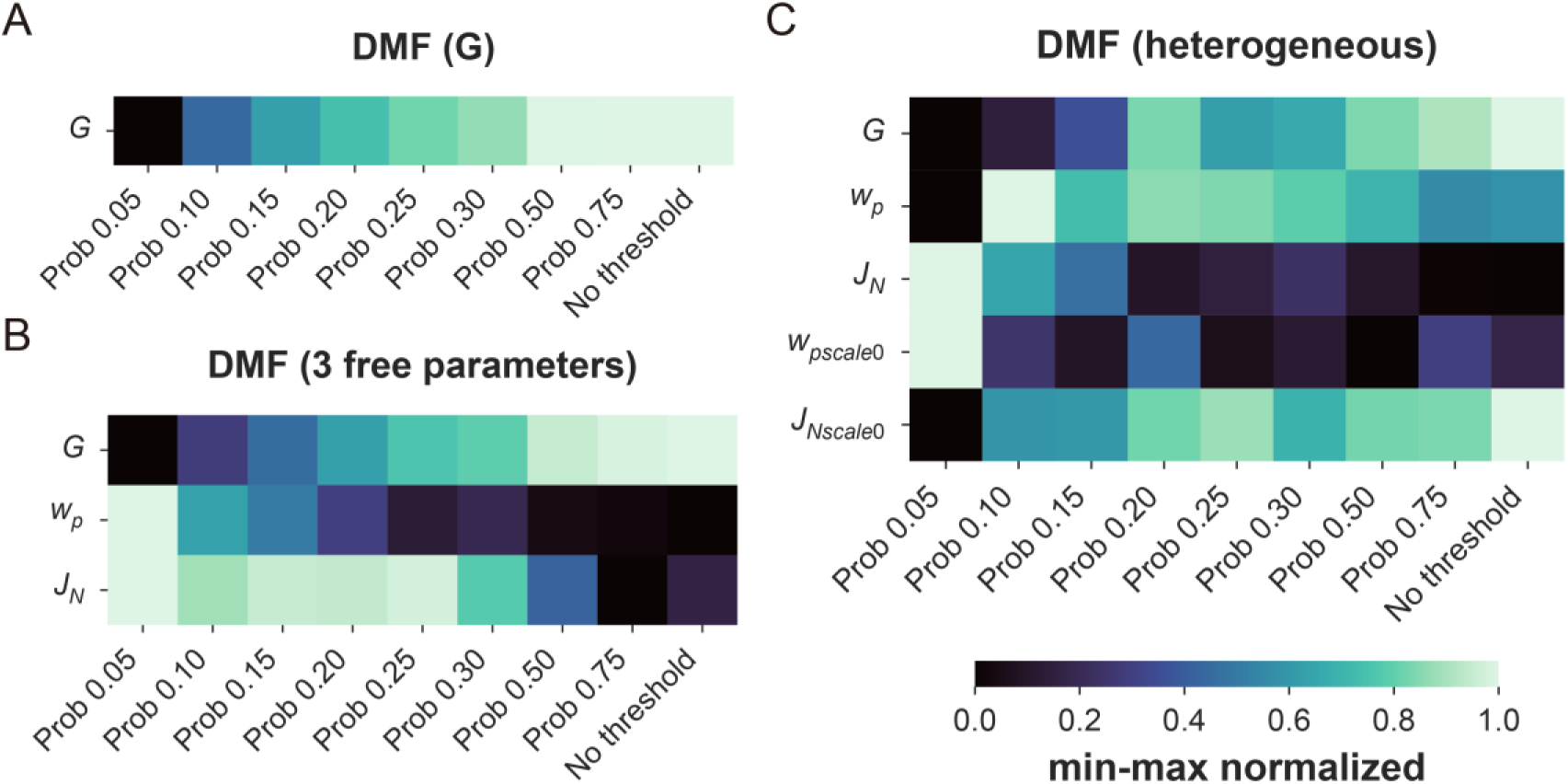
Systematic shifts in optimal biophysical parameters across varying SC densities. (A) Min-max normalized optimal global coupling parameter (*G*) for the single-parameter DMF model across different proportional thresholding densities. (B) Min-max normalized optimal parameters for the extended DMF model with three free parameters: global coupling (*G*), recurrent excitatory connection weight (*w_p_*), and NMDA receptor conductance (*J_N_*). (C) Min-max normalized optimal parameters for the heterogeneous DMF model, including baseline parameters (*G*, *w_p_*, *J_N_*) and the scaling factors for regional heterogeneity (*w_pscale_*_0_, *J_Nscale_*_0_). The color gradient represents the min-max normalized parameter values ranging from 0.0 to 1.0. Data were presented as mean values across subjects.

Furthermore, when regional heterogeneity was introduced, the baseline values for *w_p_* increased, while the scaling of the heterogeneity map decreased with rising density. *J_N_* exhibited an opposing pattern of change under these conditions. Simultaneously, *G* continued to demonstrate a consistent pattern of rising with network density (Fig. 4C).

### 3.5. Effects of structural sparsity across multiple spatial brain atlases

Recognizing that network density is inherently coupled with parcellation granularity, we further evaluated the impact of SC density on model performance across various spatial scales to broaden the scope of our benchmarking (Fig. 4A-D). We tested the Desikan-Killiany (DK) atlas (68 nodes, anatomically-defined), the functionally-derived Schaefer 100–300 series, and a hybrid configuration (Schaefer-100 plus 14 subcortical nodes). The structural topographies and edge-length distributions across these scales are detailed in Supplementary Fig. 3–6. Since even a 0.05 density threshold retained a substantial number of connections for the higher-resolution Schaefer 200 and 300 atlases, we implemented an additional fine-grained exploration within the low-density regime, varying density from 0.01 to 0.10 with a step size of 0.01.

Overall, the performance metrics of the DMF model gradually deteriorated as the number of nodes increased (Figure 5E, Supplementary Fig. 7). However, while finer parcellations generally required more stringent sparsification to maintain stability, the influence of SC density on model performance followed strikingly divergent trajectories across different spatial scales (Fig. 5F–H).

**Fig. 5.**
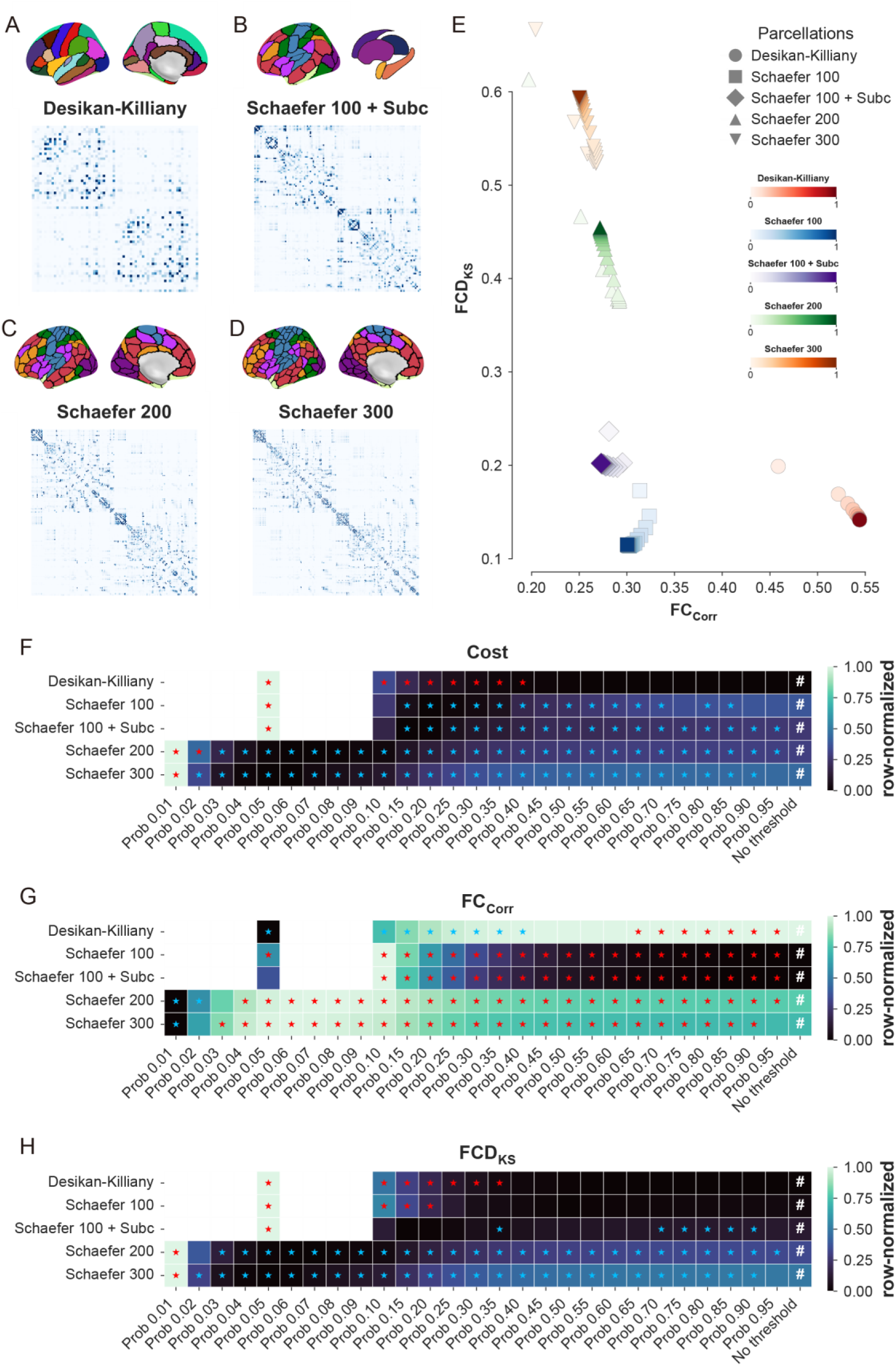
Impact of SC density on model performance across varying spatial scales. (A-D) Brain parcellation schemes and their representative structural connectivity matrices for the anatomically-defined Desikan-Killiany atlas (A), the hybrid Schaefer 100 + Subcortical regions configuration (B), Schaefer 200 (C), and Schaefer 300 (D). (E) *FC_Corr_* and *FCD_KS_* across different parcellations and SC densities, color intensity represents density, while shapes indicate parcellation types. (F-H) Evaluation of model fitting across different parcellation granularities and network densities, quantified by the row-normalized (min-max scaled) integrated Cost metric (F), *FC_Corr_* (G), and *FCD_KS_* (H). The ’#’ symbol denotes the unthresholded baseline. Stars indicate statistically significant differences compared to the baseline, blue indicates a significant decrease, while red indicates a significant increase (*P* < 0.05, paired t-test). Data were presented as mean values across subjects. Refer to Supplementary Fig. 7 for the non-normalized, raw performance metrics values.

In the low-resolution DK model, nearly all performance indices declined sharply when SC density fell to 0.40 or below; conversely, retaining the unthresholded SC yielded a relatively satisfactory performance. In contrast, for the Schaefer 100 parcellation (regardless of subcortical inclusion), moderate thresholding facilitated improvements in *FC_Corr_* without compromising the fit of *FCD_KS_*.

In contrast to the divergent patterns observed at coarser scales, the higher-resolution Schaefer 200 and 300 parcellations exhibited a concurrent optimization of multiple performance metrics during sparsification. In these fine-grained configurations, reducing SC density not only enhanced the static functional fit (*FC_Corr_*) but also significantly improved the dynamic *FCD_KS_* fit simultaneously. Except for the extreme sparsity level of 0.01—where the emergence of disconnected nodes may impede network-wide dynamics—all three performance metrics benefited from both moderate and stringent thresholding. For the Schaefer 200 atlas, the optimal model performance was localized at a density of approximately 0.07–0.08. As the parcellation granularity further increased in the Schaefer 300 atlas, this optimal performance window exhibited a downward shift toward a sparser range of approximately 0.06–0.07.

### 3.6. Correlation between structural network topology and model performance

To investigate whether the parcellation- and density-dependent variations in model performance could be attributed to the topological characteristics of the SC networks, we tracked standard graph-theoretical metrics—global efficiency ( *E_glob_*), clustering coefficient ( *C*), modularity ( *Q*), and small-worldness propensity (**φ**)—across all sparsity levels and parcellation schemes.

The trajectories of SC topological metrics across different parcellations and densities were related to, yet distinct from, those of the DMF model (Fig. 6). In contrast to the complex, non-monotonic performance patterns observed in the DMF model, the topological metrics predominantly exhibited monotonic changes. The optimal densities for *E_glob_*, *C*, and *Q* shifted toward lower values as parcellation granularity increased. Notably, these topological optima did not align with the optimal densities for DMF model performance, generally occurring at sparser levels. Conversely, small-worldness ( **φ**) consistently peaked at higher, denser configurations.

**Fig. 6.**
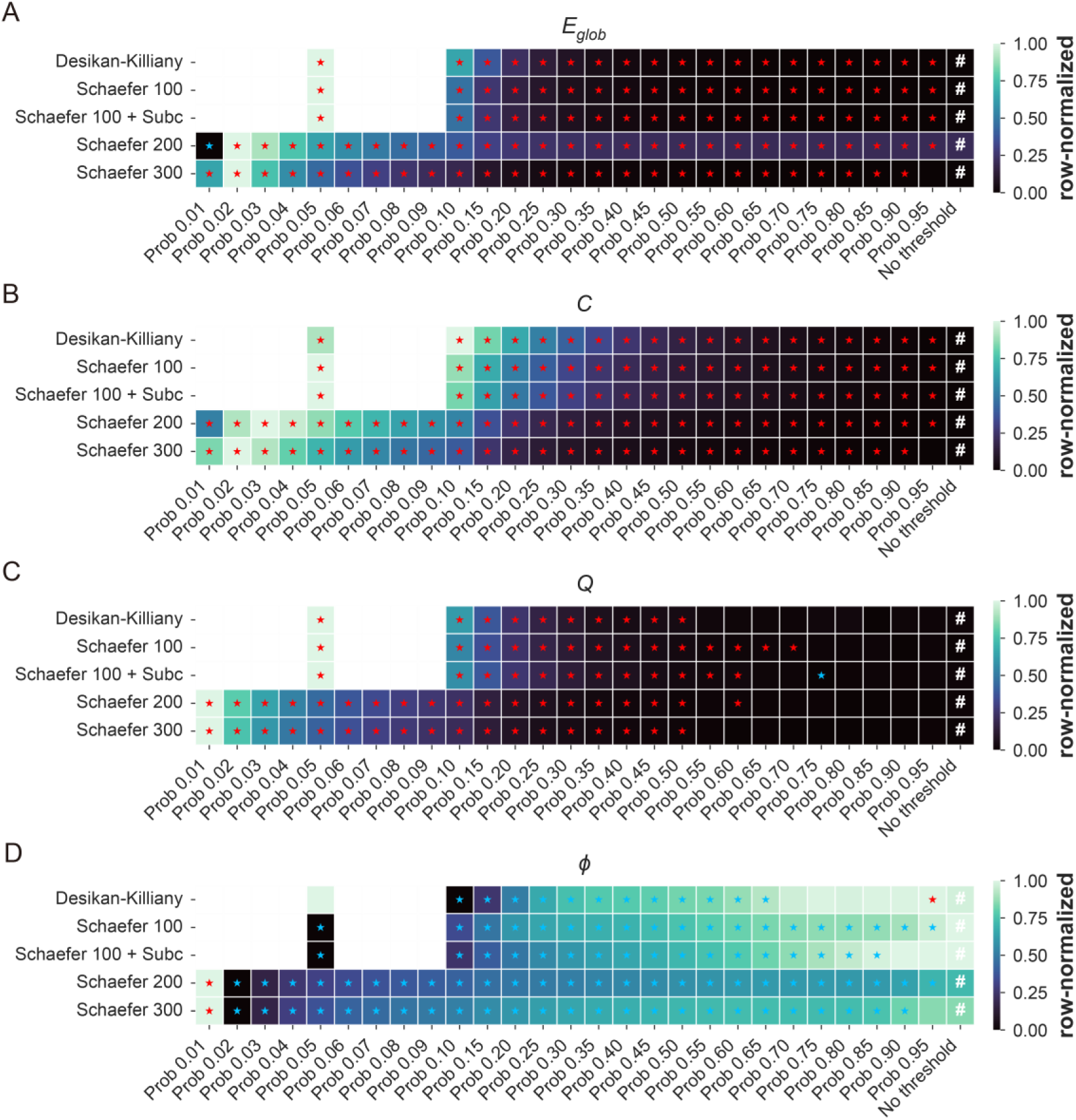
Trajectories of SC topological metrics across varying spatial scales and network densities. (A-D) Evaluation of graph-theoretical topological metrics for SC networks across different parcellation granularities and densities, quantified by the row-normalized global efficiency *E_glob_* (A), clustering coefficient C (B), modularity Q (C), and small-worldness propensity φ (D). The ’#’ symbol denotes the unthresholded baseline. Stars indicate statistically significant differences compared to the baseline, blue indicates a significant decrease, while red indicates a significant increase (*P* < 0.05, paired t-test). Data were presented as mean values across subjects. Refer to Supplementary Fig. 8 for the non-normalized, raw metrics values.

To further quantify these relationships, we computed the Spearman rank correlations between the SC topological metrics and DMF performance indices across all densities for each subject and parcellation scheme (Fig. 7). The associations between model performance and SC topology varied significantly across parcellations, occasionally reversing direction entirely. For instance, in the DK atlas, overall model performance (*Cost*, *FCD_KS_*, and *FC_Corr_*) improved in SC networks with higher *E_glob_*, *Q*, and *C*. In contrast, the high-resolution Schaefer 200 and 300 parcellations exhibited the exact opposite trend.

**Fig. 7.**
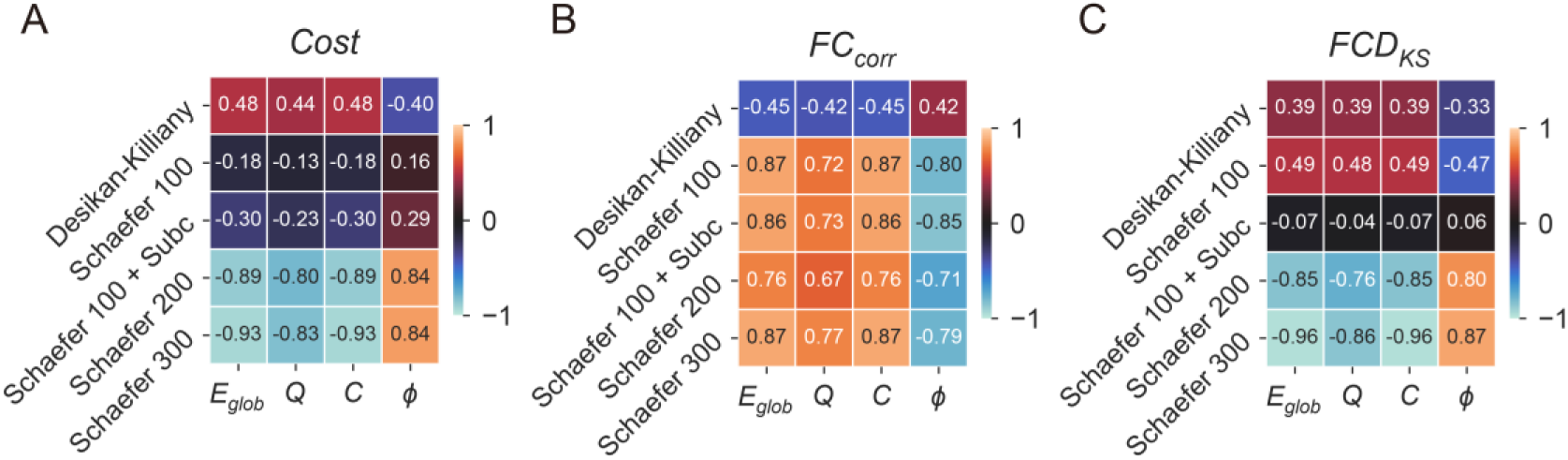
Associations between SC topology and DMF performance across spatial scales. (A-C) Spearman rank correlation coefficients between Structural Connectivity (SC) topological metrics (global efficiency *E_glob_*, modularity *Q*, clustering coefficient *C*, and small-worldness propensity φ and DMF model performance indices across different parcellation schemes and density levels. The heatmaps illustrate the correlations for the integrated Cost metric (A), *FC_corr_* (B), and *FCD_KS_* (C). The color gradients and annotated values within each cell represent the average Spearman correlation coefficient across subjects.

Specifically, for the static *FC_Corr_*, the correlation patterns with SC topology were highly consistent across all functionally defined Schaefer atlases, whereas these associations were notably weaker in the anatomically defined DK atlas.

For the dynamic *FCD_KS_*, the relationships were considerably more complex: in coarse-grained parcellations, *FCD_KS_* correlated positively with *E_glob_*, *Q*, and *C*, but negatively with **φ**. Conversely, fine-grained parcellations displayed a complete inversion of this correlational pattern.

## 4. Discussion

In the present study, we systematically investigated the impact of SC thresholding on the performance of individualized whole-brain network models. By benchmarking various thresholding strategies across diverse spatial scales, model architectures, and clinical conditions, we demonstrated that the optimal structural sparsity is fundamentally resolution-dependent. Specifically, while coarse-grained parcellations require denser connectomes to maintain accurate FCD alongside improved static FC, fine-grained models distinctly benefit from stringent proportional thresholding to simultaneously optimize both static and dynamic fits. Our findings challenge the conservative, uncritical use of raw structural connectomes and establish resolution-tailored proportional thresholding as an indispensable methodological step for robust precision brain network modelling.

### 4.1 The necessity and methodology of connectome thresholding in whole-brain modelling

Our study builds upon lessons from previous research on SC thresholding. Within the domain of graph theoretical analysis, the necessity of SC preprocessing and the choice of specific methods have long been subjects of debate. For instance, evidence suggests that the removal of weak connections may not significantly alter common global network metrics, and excessive sparsification can even introduce spurious variation (Civier et al., 2019), particularly when utilizing state-of-the-art tractography filtering techniques (Frigo et al., 2020). Conversely, other studies have demonstrated that optimizing the SC scaffold through appropriate thresholding strategies not only enhances the reliability of complex topological analyses such as community and hub identification (Dimitriadis et al., 2021) but also improves the test-retest reliability of the SC itself (Messaritaki et al., 2019). Therefore, a single strategy may exert markedly different impact patterns depending on the specific metrics being evaluated.

This phenomenon is explicitly reflected in our modelling results. In our analysis using the Schaefer-100 parcellation, which represents one of the most common spatial scales in current whole-brain modelling studies, SC thresholding revealed divergent effects on FC and FCD. FC typically benefited from thresholding, especially at relatively sparse densities. In contrast, FCD performance deteriorated significantly under such high sparsity and only achieved satisfactory fits when the SC density exceeded a certain level. This suggests that thresholding is largely optional for the FCD metric at this scale. It is important to note that this trend is not limited to individualized modelling based on subject-specific SC but also applies to individualized modelling utilizing group-level SC templates, which are commonly employed in clinical studies where high-quality individual dMRI data are unavailable (Idesis et al., 2024; Schulte et al., 2026). This pattern was further corroborated in our independent stroke dataset, emphasizing the value of benchmarking thresholding strategies in real-world clinical scenarios, even in complex cases where the brain contains structural lesions.

Notably, while the integrated Cost metric in the healthy cohort reached a global minimum at a density of 0.20, where the gains in FC fit outweighed the losses in FCD fit, this distinct minimum was not observed in the stroke cohort, nor did it emerge in our models incorporating spatial heterogeneity. Because a universal minimum that simultaneously satisfies the requirements of both FC and FCD is not always present, rather than focusing on such heuristics designed primarily for model fitting, a more robust strategy for researchers is to weigh the required thresholding density based on whether their specific simulation objectives prioritize static or dynamic functional architecture.

Furthermore, we found that highly conservative strategies often used in previous modelling studies, such as applying a minimal absolute cutoff for streamline counts (Rocha et al., 2022), offered only negligible improvements compared to an appropriate and equally simple proportional strategy. Interestingly, the OMST method, which is favoured in graph theoretical studies for its ability to preserve core topological structures without manual thresholding (Dimitriadis et al., 2017, 2021; Messaritaki et al., 2019), led to a significant decline across all DMF model performance metrics. This suggests that conclusions derived from structural graph topology cannot be directly extrapolated to whole-brain dynamic simulations, a discrepancy we explore in detail in the following sections.

### 4.2 Density effects persist across model complexities and architectures

In our primary analysis, we optimized a single free parameter, *G*, following the classic strategy of reproducing rich brain functional patterns with minimal parameters (Deco, Ponce-Alvarez, et al., 2014) while reducing the computational cost of the benchmark. Recent studies, however, have incorporated additional degrees of freedom to achieve superior fits (Deco et al., 2018, 2021; Kong et al., 2021; Saberi, Wischnewski, et al., 2025; P. Wang et al., 2019), which is particularly vital for individualized simulations. Unlike group-level models focusing on shared macroscopic mechanisms, personalized models require a framework that more closely mirrors an individual’s specific brain architecture to enable precise inference and virtual intervention.

Our results demonstrate that the impact of SC density on model performance is largely independent of parameter complexity. Whether using the three-parameter DMF model or the increasingly popular spatially heterogeneous models, the dependency of FC and FCD on network density remains consistent. Remarkably, this effect persists even when transitioning to the Kuramoto model, which operates on fundamentally different mathematical principles than the DMF model. These consistent findings likely reflect a fundamental trade-off in SC thresholding: while sparsification removes false-positive connections to highlight the structural backbone, it inevitably prunes genuine long-range connections. Previous research has underscored that these long-range projections are indispensable for generating complex temporal dynamics (Vohryzek et al., 2025), explaining why FCD performance suffers as networks become overly sparse.

A primary utility of high-degree-of-freedom individualized models is the use of inferred parameters as personalized features or biomarkers (Deco et al., 2026; Schulte et al., 2026), however, we found that SC thresholding strategies induce systematic shifts not only in model performance but also in the resulting biophysical parameter estimates. At higher SC densities, the model requires a higher *G* to reach the optimal dynamical regime, while the inferred values for *w_p_* and *J_N_* concurrently decrease. Interestingly, network density even modulates how the model utilizes regional heterogeneity maps. Under high-density conditions, the scaling of *w_p_* by the myelin map decreases, whereas the scaling of *J_N_* increases. These systematic shifts suggest that any neurobiological interpretation of these parameters must be handled with extreme caution. We emphasize that when treating these biophysical parameters as individual traits, researchers should explicitly report the density and thresholding characteristics of the underlying SC to ensure valid comparisons across different studies.

### 4.3 The interplay between network resolution and structural sparsity

SC density cannot be discussed in isolation from the underlying parcellation scheme, as finer granularities inherently lead to a higher proportion of disconnected nodes. Our study systematically evaluated a diverse range of spatial scales, from coarse-grained DK and Schaefer-100 to high-resolution Schaefer-200 and 300 parcellations. The latter, though less common, are reported to yield the richest dynamic repertoire (Kobeleva et al., 2021). We also incorporated subcortical nodes into the Schaefer-100 framework, recognizing that subcortical inclusion is essential for clinical modelling needs such as deep brain stimulation (An et al., 2022; Zhang et al., 2025).

Our findings reveal substantial heterogeneity in how different parcellations respond to sparsification. For the most coarse-grained DK atlas, the unthresholded SC yielded near-optimal performance. This is in stark contrast to the Schaefer-100 results, and the discrepancy became even more pronounced at higher resolutions. For the Schaefer-200 and Schaefer-300 atlases, almost any degree of thresholding led to a systematic improvement in all performance metrics. Crucially, this improvement extended to FCD, which, unlike in coarse-grained parcellations, did not suffer from moderate or even stringent sparsification.

These resolution-dependent patterns are likely rooted in the inherent limitations of diffusion tractography. Previous research mapping the human connectome has demonstrated that even high-quality imaging data can yield over 60% false-positive fibres (Maier-Hein et al., 2017). Direct comparisons between diffusion MRI tractography and gold-standard anatomical tracing in macaques have reached similar conclusions (Thomas et al., 2014), confirming that high sensitivity often comes at the cost of low specificity (Sarwar et al., 2019; Tahedl et al., 2025; C.-H. Yeh et al., 2021). This provides a plausible explanation for the observed results: in coarse-grained parcellations, the regions are large enough that very few inter-regional edges are composed entirely of false positives. At this scale, thresholding struggles to distinguish between true and false components within a single edge; thus, pruning primarily risks deleting genuine anatomical connections. In contrast, as parcellation granularity increases, tractography continues to produce nearly dense matrices, but a much larger proportion of these fine-grained edges represent purely spurious connections that do not exist anatomically. In this high-resolution regime, thresholding effectively prunes these weak, false-positive "short circuits" without significantly compromising the core long-range backbone.

### 4.4 Connectome topology does not fully explain dynamical modelling optima

The discovery that model performance depends on both SC density and parcellation granularity presents a significant challenge for researchers using alternative spatial scales, as no universal thresholding parameter appears applicable across all levels of resolution. A potential solution would be to utilize computationally efficient graph theoretical metrics that are also sensitive to SC structure (Corso et al., 2026; Zalesky et al., 2010) to pre-select an optimal thresholding strategy for a given parcellation.

Our results demonstrate that this approach is currently unfeasible for guiding thresholding choices. While previous literature has indicated that topological metrics often fail to explain individual variations in model performance across different parcellations (Domhof et al., 2021), we replicated and extended this finding by systematically analysing classical graph metrics across multiple node granularities and densities. Although graph metrics exhibit certain patterns consistent with our modelling results—such as finer parcellations requiring more stringent thresholding to achieve optimal topological properties while coarse parcellations remain relatively insensitive—the specific trajectories and optimal peaks of these metrics diverge significantly from the actual DMF performance indices. This divergence is particularly pronounced when independently considering the fits for FC and FCD. The lack of alignment was further confirmed by the inconsistent correlations between individual model fitting curves and topological metric curves across the density spectrum. Remarkably, the same graph metric could demonstrate a positive correlation with model performance in one parcellation while showing a negative correlation in another. These inconsistencies suggest that topological properties derived purely from SC are insufficient for explaining or predicting the behaviour of biophysical models like the DMF.

This disconnect likely explains why algorithms designed specifically for graph theoretical analysis, such as OMST, perform poorly in the context of dynamical modelling. Unlike pure structural analysis, whole-brain modelling represents a complex SC-FC coupling problem (Turnbull et al., 2025). While also sensitive to SC, the interplay between the anatomical scaffold and neuronal dynamics can introduce non-linearities beyond the reach of simple topological metrics.

Given the inability to derive reliable prior knowledge from SC topology to support the selection of an optimal threshold and considering the variations introduced by imaging quality and tractography algorithms, we propose a pragmatic approach. Until more advanced thresholding strategies emerge that can accurately infer true anatomical connectivity without manual parameter tuning, it is essential for researchers to conduct small-sample benchmark tests to identify the optimal threshold. This step is particularly indispensable for large-scale studies that prioritize individualized fitting or utilize fine-grained parcellations.

### 4.5 Methodological recommendations for future studies

Based on our benchmarking results, we propose the following actionable guidelines for future personalized whole-brain modelling studies:

1. Avoid purely topology-driven thresholding: Methods designed to optimize graph-theoretical properties (e.g., OMST) are suboptimal for biophysical generative models.
2. Tailor thresholds to the target metric in coarse parcellations: For atlases like Schaefer-100, researchers must prioritize. If the study focuses exclusively on FCD, raw SCs are acceptable. If simulating precise static FC is the goal, moderate proportional thresholding is recommended.
3. Always threshold high-resolution models: For fine-grained parcellations (e.g., ≥ 200 nodes), strict proportional thresholding is universally beneficial.
4. Conduct preliminary benchmarking: Our recommendations are tailored to the methodologies described herein. For studies that deviate significantly in experimental protocols or data characteristics, we strongly advise conducting an empirical benchmark sweep on a small sample set. This ensures the threshold is calibrated before proceeding with computationally intensive, large-scale simulations.

### 4.6 Limitations and future directions

Several limitations of the current study must be acknowledged.

First, our structural connectomes were derived from multi-shell data using a probabilistic dMRI pipeline with high streamline counts setting. However, because the resulting SC is highly sensitive to acquisition (Jones, 2004), preprocessing (Seider et al., 2022), and the total number of streamlines tracked (Jung et al., 2021), our findings must be interpreted within the context of the specific workflow. Furthermore, deterministic tractography implemented in tools like DSI Studio (F.-C. Yeh, 2025) inherently produces distinct sparsity profiles. This variability suggests that further comparative investigation is required.

Second, while our benchmark systematically accounts for factors such as parcellation granularity, model architectures, and parameter complexity, it does not claim to be an exhaustive exploration of all possible configurations. For instance, our analysis of high-resolution parcellations was specifically restricted to the optimization of a single global coupling coefficient. These simplifications, while necessary for computational tractability, mean that the interaction between network density and more complex physiological constraints remains to be fully elucidated.

Third, we only utilized a group-level SC matrix derived from the median across HCP subjects to evaluate how network density impacts the performance of group-level SC-based DMF models. Given the multitude of consensus- averaging algorithms available and the continued importance of group-level templates in datasets lacking individual SC, the interaction between specific consensus methodologies and density thresholds represents a critical avenue for future exploration.

Fourth, our investigation was restricted to the DMF and Kuramoto models and focused exclusively on BOLD-derived signals. There are many other widely used biophysical and phenomenological models, as well as frameworks designed to infer electrophysiological activity such as EEG or MEG (Breakspear, 2017). Future research should aim to incorporate a broader range of model architectures and signal modalities to further validate the generalizability of these results across different neural imaging scales.

Finally, there are studies that do not rely solely on raw structural connectomes. Instead, they first optimize the SC matrix by adding anatomical links and reweighting edges to maximize structure–function interplay (Deco, McIntosh, et al., 2014). Others employ the Hopf model to infer effective connectivity (Deco et al., 2019). Our current benchmark did not evaluate these inferential optimizations, focusing instead on the direct implementation of structural scaffolds.

Looking forward, future methodological efforts should focus on designing bespoke SC thresholding algorithms optimized specifically for whole-brain modelling—methods that successfully prune false-positive tractography noise while faithfully preserving the underlying connectivity weights essential for driving realistic large-scale neural dynamics.

## 5 Data and code availability

The MICA-MICs and HCP-YA datasets are openly available via the Canadian Open Neuroscience Platform (https://db.humanconnectome.org/) and the BALSA (https://balsa.wustl.edu/), respectively. Clinical data from post-stroke patients are not publicly available due to ethical restrictions but may be shared by the corresponding authors upon reasonable request.

All simulations were implemented using the cuBNM framework (https://github.com/amnsbr/cubnm). Custom scripts for model configuration and parameter optimization are available on GitHub at https://github.com/Qian-Liyi/BNM_thrSC.

## 6 Author contributions

LQ and SJ contributed equally to this work. LQ: Conceptualization, Methodology, Software, Formal analysis, Visualization, Writing – original draft. SJ: Conceptualization, Methodology, Investigation, Resources, Writing – original draft. ZL: Investigation, Data curation. YX: Resources, Investigation. QY: Validation, Supervision, Writing – review & editing. LC: Conceptualization, Supervision, Project administration, Funding acquisition, Writing – review & editing.

