## Supplementary methods and figures for "Systematic evaluation of structural connectome thresholding in whole-brain network modelling"

**Affiliations:**

### Supplementary Methods S1.

Preprocessing was performed using *QSIPrep* 1.0.1.dev0+gee9aa2e.d20250115 (Cieslak et al. 2021), which is based on *Nipype* 1.9.1 [Gorgolewski et al. (2011); Gorgolewski et al. (2018); RRID:SCR_002502].

The anatomical reference image was reoriented into AC-PC alignment via a 6-DOF transform extracted from a full Affine registration to the MNI152NLin2009cAsym template. A full nonlinear registration to the template from AC-PC space was estimated via symmetric nonlinear registration (SyN) using antsRegistration. Brain extraction was performed on the T1w image using SynthStrip (Hoopes et al. 2022) and automated segmentation was performed using SynthSeg (Billot, Greve, et al. 2023, @synthseg2) from FreeSurfer version 7.3.1.

#### Diffusion data preprocessing

DWI data were denoised using the Marchenko-Pastur PCA method implemented in dwidenoise (Tournier et al. 2019; Veraart et al. 2016; Veraart, Fieremans, and Novikov 2016; Cordero-Grande et al. 2019) with an automatically-determined window size of 3 voxels. Any images with a b-value less than 100 s/mm^2 were treated as a *b*=0 image. The mean intensity of the DWI series was adjusted so all the mean intensity of the b=0 images matched across eachseparate DWI scanning sequence. B1 field inhomogeneity was corrected using dwibiascorrect from MRtrix3 with the N4 algorithm (Tustison et al. 2010) after corrected images were resampled.

FSL (version None)’s eddy was used for head motion correction and Eddy current correction (Andersson and Sotiropoulos 2016). Eddy was configured with a *q*-space smoothing factor of 10, a total of 5 iterations, and 1000 voxels used to estimate hyperparameters. A quadratic first level model and a linear second level model were used to characterize Eddy current-related spatial distortion. *q*-space coordinates were forcefully assigned to shells. Field offset was attempted to be separated from subject movement. Shells were aligned post-eddy. Eddy’s outlier replacement was run (Andersson et al. 2016). Data were grouped by slice, only including values from slices determined to contain at least 250 intracerebral voxels. Groups deviating by more than 4 standard deviations from the prediction had their data replaced with imputed values. Final interpolation was performed using the jac method.

Several confounding time-series were calculated based on the preprocessed DWI: framewise displacement (FD) using the implementation in *Nipype* (following the definitions by Power et al. 2014). The head-motion estimates calculated in the correction step were also placed within the corresponding confounds file. Slicewise cross correlation was also calculated. The DWI time-series were resampled to ACPC, generating a *preprocessed DWI run in ACPC space* with 1.5mm isotropic voxels.

Many internal operations of *QSIPrep* use *Nilearn* 0.10.1 (Abraham et al. 2014, RRID:SCR_001362) and *Dipy* (Garyfallidis et al. 2014). For more details of the pipeline, see [the section corresponding to workflows in *QSIPrep*’s documentation](https://qsiprep.readthedocs.io/en/latest/workflows.html).

Billot, Benjamin, Douglas N Greve, Oula Puonti, Axel Thielscher, Koen Van Leemput, Bruce Fischl, Adrian V Dalca, Juan Eugenio Iglesias, and others. 2023. “SynthSeg: Segmentation of Brain Mri Scans of Any Contrast and Resolution Without Retraining.” *Medical Image Analysis* 86. Elsevier: 102789.

Billot, Benjamin, Colin Magdamo, You Cheng, Steven E Arnold, Sudeshna Das, and Juan Eugenio Iglesias. 2023. “Robust Machine Learning Segmentation for Large-Scale Analysis of Heterogeneous Clinical Brain Mri Datasets.” *Proceedings of the National Academy of Sciences* 120 (9). National Acad Sciences: e2216399120.

Cieslak, Matthew, Philip A Cook, Xiaosong He, Fang-Cheng Yeh, Thijs Dhollander, Azeez Adebimpe, Geoffrey K Aguirre, et al. 2021. “QSIPrep: An Integrative Platform for Preprocessing and Reconstructing Diffusion Mri Data.” *Nature Methods* 18 (7). Nature Publishing Group US New York: 775–78. <https://doi.org/10.1038/s41592-021-01185-5>.

Cordero-Grande, Lucilio, Daan Christiaens, Jana Hutter, Anthony N Price, and Jo V Hajnal. 2019. “Complex Diffusion-Weighted Image Estimation via Matrix Recovery Under General Noise Models.” *Neuroimage* 200. Elsevier: 391–404. <https://doi.org/10.1016/j.neuroimage.2019.06.039>.

Garyfallidis, Eleftherios, Matthew Brett, Bagrat Amirbekian, Ariel Rokem, Stefan Van Der Walt, Maxime Descoteaux, and Ian Nimmo-Smith. 2014. “Dipy, a Library for the Analysis of Diffusion Mri Data.” *Frontiers in Neuroinformatics* 8. Frontiers: 8.

Gorgolewski, K., C. D. Burns, C. Madison, D. Clark, Y. O. Halchenko, M. L. Waskom, and S. Ghosh. 2011. “Nipype: A Flexible, Lightweight and Extensible Neuroimaging Data Processing Framework in Python.” *Frontiers in Neuroinformatics* 5: 13. <https://doi.org/10.3389/fninf.2011.00013>.

Gorgolewski, Krzysztof J., Oscar Esteban, Christopher J. Markiewicz, Erik Ziegler, David Gage Ellis, Michael Philipp Notter, Dorota Jarecka, et al. 2018. “Nipype.” *Software*. Zenodo. <https://doi.org/10.5281/zenodo.596855>.

Hoopes, Andrew, Jocelyn S Mora, Adrian V Dalca, Bruce Fischl, and Malte Hoffmann. 2022. “SynthStrip: Skull-Stripping for Any Brain Image.” *NeuroImage* 260. Elsevier: 119474.

Power, Jonathan D., Anish Mitra, Timothy O. Laumann, Abraham Z. Snyder, Bradley L. Schlaggar, and Steven E. Petersen. 2014. “Methods to Detect, Characterize, and Remove Motion Artifact in Resting State fMRI.” *NeuroImage* 84 (Supplement C): 320–41. <https://doi.org/10.1016/j.neuroimage.2013.08.048>.

Tournier, J-Donald, Robert Smith, David Raffelt, Rami Tabbara, Thijs Dhollander, Maximilian Pietsch, Daan Christiaens, Ben Jeurissen, Chun-Hung Yeh, and Alan Connelly. 2019. “MRtrix3: A Fast, Flexible and Open Software Framework for Medical Image Processing and Visualisation.” *Neuroimage* 202. Elsevier: 116137. <https://doi.org/10.1016/j.neuroimage.2019.116137>.

Tustison, N. J., B. B. Avants, P. A. Cook, Y. Zheng, A. Egan, P. A. Yushkevich, and J. C. Gee. 2010. “N4ITK: Improved N3 Bias Correction.” *IEEE Transactions on Medical Imaging* 29 (6): 1310–20. <https://doi.org/10.1109/TMI.2010.2046908>.

Veraart, Jelle, Els Fieremans, and Dmitry S Novikov. 2016. “Diffusion Mri Noise Mapping Using Random Matrix Theory.” *Magnetic Resonance in Medicine* 76 (5). Wiley Online Library: 1582–93.

Veraart, Jelle, Dmitry S Novikov, Daan Christiaens, Benjamin Ades-Aron, Jan Sijbers, and Els Fieremans. 2016. “Denoising of Diffusion Mri Using Random Matrix Theory.” *NeuroImage* 142. Elsevier: 394–406.

### Supplementary Methods S2

Reconstruction was performed using *QSIRecon* 1.0.1.dev0+gf78c888.d20250114 (Cieslak et al. (2021)), which is based on *Nipype* 1.9.1 (Gorgolewski et al. (2011); Gorgolewski et al. (2018); RRID:SCR_002502).

A hybrid surface/volume segmentation was created [Smith 2020].FreeSurfer outputs were registered to the QSIRecon outputs.

#### Anatomical data for DWI reconstruction

T1w-based spatial normalization calculated during preprocessing was used to map atlases from template space into alignment with DWIs. Brainmasks from antsBrainExtraction were used in all subsequent reconstruction steps. The following atlases were used in the workflow: the Schaefer Supplemented with Subcortical Structures (4S) atlas (Schaefer et al. 2018; Pauli, Nili, and Tyszka 2018; King et al. 2019; Najdenovska et al. 2018; Glasser et al. 2013) at 3 different resolutions (156, 256, and 456 parcels), the Automated Anatomical Labeling (AAL) 116-parcel atlas (Tzourio-Mazoyer et al. 2002), the AICHA 384-parcel atlas (Joliot et al. 2015) extended with subcortical parcels, the Brainnetome 246-parcel atlas (Fan et al. 2016) extended with subcortical parcels, and the Gordon 333-parcel atlas (Gordon et al. 2016) extended with subcortical parcels. Cortical parcellations were mapped from template space to DWIS using the T1w-based spatial normalization.

#### MRtrix3 Reconstruction

Multi-tissue fiber response functions were estimated using the dhollander algorithm. FODs were estimated via constrained spherical deconvolution (CSD, Tournier et al. (2004), Tournier et al. (2008)) using an unsupervised multi-tissue method (Dhollander et al. (2019), Dhollander, Raffelt, and Connelly (2016)). Reconstruction was done using MRtrix3 (J-Donald et al. (2019)). FODs were intensity-normalized using mtnormalize (Raffelt et al. (2017)).

Many internal operations of *QSIRecon* use *Nilearn* 0.10.1 (Abraham et al. 2014, RRID:SCR_001362) and *Dipy* 1.8.0(Garyfallidis et al. 2014). For more details of the pipeline, see [the section corresponding to workflows in *QSIRecon*’s documentation](https://qsirecon.readthedocs.io/en/latest/workflows.html).

Garyfallidis, Eleftherios, Matthew Brett, Bagrat Amirbekian, Ariel Rokem, Stefan Van Der Walt, Maxime Descoteaux, and Ian Nimmo-Smith. 2014. “Dipy, a Library for the Analysis of Diffusion Mri Data.” *Frontiers in Neuroinformatics* 8. Frontiers: 8.

Glasser, Matthew F., Stamatios N. Sotiropoulos, J. Anthony Wilson, Timothy S. Coalson, Bruce Fischl, Jesper L. Andersson, Junqian Xu, et al. 2013. “The Minimal Preprocessing Pipelines for the Human Connectome Project.” *NeuroImage*, Mapping the connectome, 80: 105–24. <https://doi.org/10.1016/j.neuroimage.2013.04.127>.

Gordon, Evan M, Timothy O Laumann, Babatunde Adeyemo, Jeremy F Huckins, William M Kelley, and Steven E Petersen. 2016. “Generation and Evaluation of a Cortical Area Parcellation from Resting-State Correlations.” *Cerebral Cortex* 26 (1). Oxford University Press: 288–303. <https://doi.org/10.1093/cercor/bhu239>.

Gorgolewski, K., C. D. Burns, C. Madison, D. Clark, Y. O. Halchenko, M. L. Waskom, and S. Ghosh. 2011. “Nipype: A Flexible, Lightweight and Extensible Neuroimaging Data Processing Framework in Python.” *Frontiers in Neuroinformatics* 5: 13. <https://doi.org/10.3389/fninf.2011.00013>.

Gorgolewski, Krzysztof J., Oscar Esteban, Christopher J. Markiewicz, Erik Ziegler, David Gage Ellis, Michael Philipp Notter, Dorota Jarecka, et al. 2018. “Nipype.” *Software*. Zenodo. <https://doi.org/10.5281/zenodo.596855>.

J-Donald, Robert Smith, David Raffelt, Rami Tabbara, Thijs Dhollander, Maximilian Pietsch, Daan Christiaens, Ben Jeurissen, Chun-Hung Yeh, and Alan" Connelly. 2019. “MRtrix3: A Fast, Flexible and Open Software Framework for Medical Image Processing and Visualisation.” *NeuroImage* 202: 116137.

Joliot, Marc, Gaël Jobard, Mikaël Naveau, Nicolas Delcroix, Laurent Petit, Laure Zago, Fabrice Crivello, Emmanuel Mellet, Bernard Mazoyer, and Nathalie Tzourio-Mazoyer. 2015. “AICHA: An Atlas of Intrinsic Connectivity of Homotopic Areas.” *Journal of Neuroscience Methods* 254. Elsevier: 46–59. <https://doi.org/10.1016/j.jneumeth.2015.07.013>.

King, Maedbh, Carlos R Hernandez-Castillo, Russell A Poldrack, Richard B Ivry, and Jörn Diedrichsen. 2019. “Functional Boundaries in the Human Cerebellum Revealed by a Multi-Domain Task Battery.” *Nature Neuroscience* 22 (8). Nature Publishing Group US New York: 1371–8. <https://doi.org/10.1038/s41593-019-0436-x>.

Najdenovska, Elena, Yasser Alemán-Gómez, Giovanni Battistella, Maxime Descoteaux, Patric Hagmann, Sebastien Jacquemont, Philippe Maeder, Jean-Philippe Thiran, Eleonora Fornari, and Meritxell Bach Cuadra. 2018. “In-Vivo Probabilistic Atlas of Human Thalamic Nuclei Based on Diffusion-Weighted Magnetic Resonance Imaging.” *Scientific Data* 5 (1). Nature Publishing Group: 1–11. <https://doi.org/10.1038/sdata.2018.270>.

Pauli, Wolfgang M, Amanda N Nili, and J Michael Tyszka. 2018. “A High-Resolution Probabilistic in Vivo Atlas of Human Subcortical Brain Nuclei.” *Scientific Data* 5 (1). Nature Publishing Group: 1–13. <https://doi.org/10.1038/sdata.2018.63>.

Raffelt, David, Thijs Dhollander, J-Donald Tournier, Rami Tabbara, Robert E Smith, Eric Pierre, and Alan Connelly. 2017. “Bias Field Correction and Intensity Normalisation for Quantitative Analysis of Apparent Fibre Density.” In *Proc. Intl. Soc. Mag. Reson. Med*, 25:3541.

Schaefer, Alexander, Ru Kong, Evan M Gordon, Timothy O Laumann, Xi-Nian Zuo, Avram J Holmes, Simon B Eickhoff, and BT Thomas Yeo. 2018. “Local-Global Parcellation of the Human Cerebral Cortex from Intrinsic Functional Connectivity Mri.” *Cerebral Cortex* 28 (9). Oxford University Press: 3095–3114. <https://doi.org/10.1093/cercor/bhx179>.

Tournier, J-Donald, Fernando Calamante, David G Gadian, and Alan Connelly. 2004. “Direct Estimation of the Fiber Orientation Density Function from Diffusion-Weighted Mri Data Using Spherical Deconvolution.” *NeuroImage* 23 (3). Elsevier: 1176–85.

Tournier, J-Donald, Chun-Hung Yeh, Fernando Calamante, Kuan-Hung Cho, Alan Connelly, and Ching-Po Lin. 2008. “Resolving Crossing Fibres Using Constrained Spherical Deconvolution: Validation Using Diffusion-Weighted Imaging Phantom Data.” *Neuroimage* 42 (2). Elsevier: 617–25.

Tzourio-Mazoyer, Nathalie, Brigitte Landeau, Dimitri Papathanassiou, Fabrice Crivello, Octave Etard, Nicolas Delcroix, Bernard Mazoyer, and Marc Joliot. 2002. “Automated Anatomical Labeling of Activations in Spm Using a Macroscopic Anatomical Parcellation of the Mni Mri Single-Subject Brain.” *Neuroimage* 15 (1). Elsevier: 273–89. <https://doi.org/10.1006/nimg.2001.0978>.

### Supplementary Methods S3

Results included in this manuscript come from preprocessing performed using *fMRIPrep* 24.1.1 (Esteban et al. (2019); Esteban et al. (2018); RRID:SCR_016216), which is based on *Nipype* 1.8.6 (K. Gorgolewski et al. (2011); K. J. Gorgolewski et al. (2018); RRID:SCR_002502).

#### Anatomical data preprocessing

A total of 1 T1-weighted (T1w) images were found within the input BIDS dataset. The T1w image was corrected for intensity non-uniformity (INU) with N4BiasFieldCorrection (Tustison et al. 2010), distributed with ANTs 2.5.3 (Avants et al. 2008, RRID:SCR_004757), and used as T1w-reference throughout the workflow. The T1w-reference was then skull-stripped with a *Nipype* implementation of the antsBrainExtraction.sh workflow (from ANTs), using OASIS30ANTs as target template. Brain tissue segmentation of cerebrospinal fluid (CSF), white-matter (WM) and gray-matter (GM) was performed on the brain-extracted T1w using fast (FSL (version unknown), RRID:SCR_002823, Zhang, Brady, and Smith 2001). Brain surfaces were reconstructed using recon-all (FreeSurfer 7.3.2, RRID:SCR_001847, Dale, Fischl, and Sereno 1999), and the brain mask estimated previously was refined with a custom variation of the method to reconcile ANTs-derived and FreeSurfer-derived segmentations of the cortical gray-matter of Mindboggle (RRID:SCR_002438, Klein et al. 2017). Volume-based spatial normalization to one standard space (MNI152NLin2009cAsym) was performed through nonlinear registration with antsRegistration (ANTs 2.5.3), using brain-extracted versions of both T1w reference and the T1w template. The following template was were selected for spatial normalization and accessed with *TemplateFlow* (24.2.0, Ciric et al. 2022): *ICBM 152 Nonlinear Asymmetrical template version 2009c* [Fonov et al. (2009), RRID:SCR_008796; TemplateFlow ID: MNI152NLin2009cAsym].

#### Functional data preprocessing

For each of the 1 BOLD runs found per subject (across all tasks and sessions), the following preprocessing was performed. First, a reference volume was generated, using a custom methodology of *fMRIPrep*, for use in head motion correction. Head-motion parameters with respect to the BOLD reference (transformation matrices, and six corresponding rotation and translation parameters) are estimated before any spatiotemporal filtering using mcflirt (FSL , Jenkinson et al. 2002). The BOLD reference was then co-registered to the T1w reference using bbregister (FreeSurfer) which implements boundary-based registration (Greve and Fischl 2009). Co-registration was configured with six degrees of freedom. Several confounding time-series were calculated based on the *preprocessed BOLD*: framewise displacement (FD), DVARS and three region-wise global signals. FD was computed using two formulations following Power (absolute sum of relative motions, Power et al. (2014)) and Jenkinson (relative root mean square displacement between affines, Jenkinson et al. (2002)). FD and DVARS are calculated for each functional run, both using their implementations in *Nipype* (following the definitions by Power et al. 2014). The three global signals are extracted within the CSF, the WM, and the whole-brain masks. Additionally, a set of physiological regressors were extracted to allow for component-based noise correction (*CompCor*, Behzadi et al. 2007). Principal components are estimated after high-pass filtering the *preprocessed BOLD* time-series (using a discrete cosine filter with 128s cut-off) for the two *CompCor* variants: temporal (tCompCor) and anatomical (aCompCor). tCompCor components are then calculated from the top 2% variable voxels within the brain mask. For aCompCor, three probabilistic masks (CSF, WM and combined CSF+WM) are generated in anatomical space. The implementation differs from that of Behzadi et al. in that instead of eroding the masks by 2 pixels on BOLD space, a mask of pixels that likely contain a volume fraction of GM is subtracted from the aCompCor masks. This mask is obtained by dilating a GM mask extracted from the FreeSurfer’s *aseg* segmentation, and it ensures components are not extracted from voxels containing a minimal fraction of GM. Finally, these masks are resampled into BOLD space and binarized by thresholding at 0.99 (as in the original implementation). Components are also calculated separately within the WM and CSF masks. For each CompCor decomposition, the *k* components with the largest singular values are retained, such that the retained components’ time series are sufficient to explain 50 percent of variance across the nuisance mask (CSF, WM, combined, or temporal). The remaining components are dropped from consideration. The head-motion estimates calculated in the correction step were also placed within the corresponding confounds file. The confound time series derived from head motion estimates and global signals were expanded with the inclusion of temporal derivatives and quadratic terms for each (Satterthwaite et al. 2013). Frames that exceeded a threshold of 0.5 mm FD or 1.5 standardized DVARS were annotated as motion outliers. Additional nuisance timeseries are calculated by means of principal components analysis of the signal found within a thin band (*crown*) of voxels around the edge of the brain, as proposed by (Patriat, Reynolds, and Birn 2017). All resamplings can be performed with *a single interpolation step* by composing all the pertinent transformations (i.e. head-motion transform matrices, susceptibility distortion correction when available, and co-registrations to anatomical and output spaces). Gridded (volumetric) resamplings were performed using nitransforms, configured with cubic B-spline interpolation.

Many internal operations of *fMRIPrep* use *Nilearn* 0.10.4 (Abraham et al. 2014, RRID:SCR_001362), mostly within the functional processing workflow. For more details of the pipeline, see [the section corresponding to workflows in *fMRIPrep*’s documentation](https://fmriprep.readthedocs.io/en/latest/workflows.html).

#### Copyright Waiver

The above boilerplate text was automatically generated by fMRIPrep with the express intention that users should copy and paste this text into their manuscripts *unchanged*. It is released under the [CC0](https://creativecommons.org/publicdomain/zero/1.0/) license.

### Supplementary Figures and Figure Legends

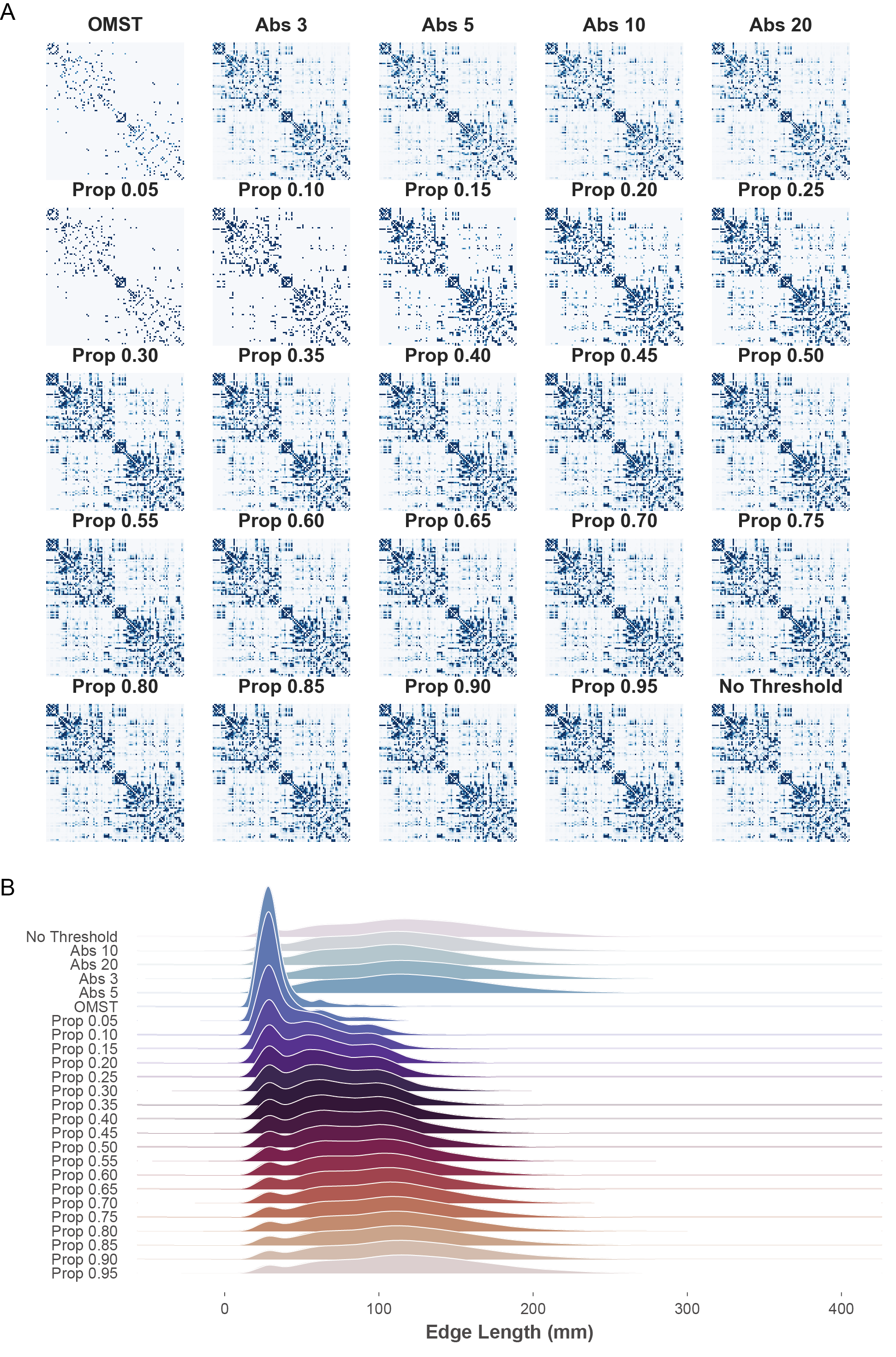

**Supplementary Fig. 1. Representative structural connectivity matrices and edge-length distributions based on the Schaefer-100.**

(A) Representative individual structural connectivity (SC) matrices based on the Schaefer-100 parcellation across varying thresholding strategies. Matrices are shown for the Orthogonal Minimum Spanning Tree (OMST), absolute streamline count thresholds (Abs) ranging from 3 to 20, proportional thresholds (Prop) ranging from 0.05 to 0.95, and the unthresholded baseline (No Threshold). (B) Density distributions of edge lengths (mm) for the connections retained under each respective thresholding strategy shown in (A).

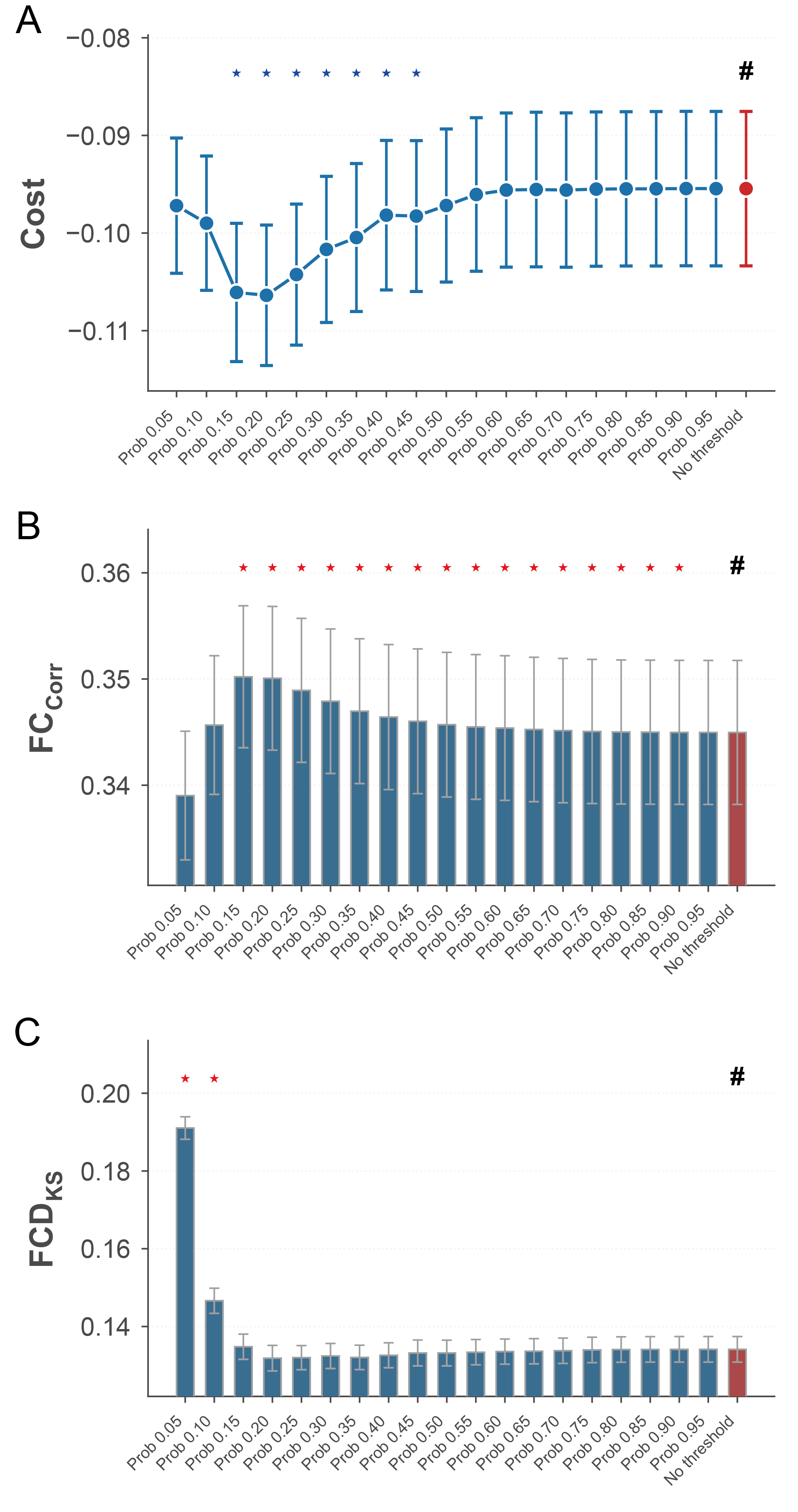

**Supplementary Fig. 2. Impact of group-level SC density on biophysical network model performance.**

(A-C) Evaluation of Dynamic Mean-Field (DMF) model fitting utilizing a consensus group-level structural connectome derived from the HCP-YA dataset. Performance is quantified by the integrated Cost metric (A), (B), and (C) across varying proportional thresholding densities. The '#' symbol denotes the unthresholded baseline. Stars indicate statistically significant differences compared to the baseline; blue indicates a significant decrease, while red indicates a significant increase (P < 0.05, paired t-test). Data are presented as mean SEM.

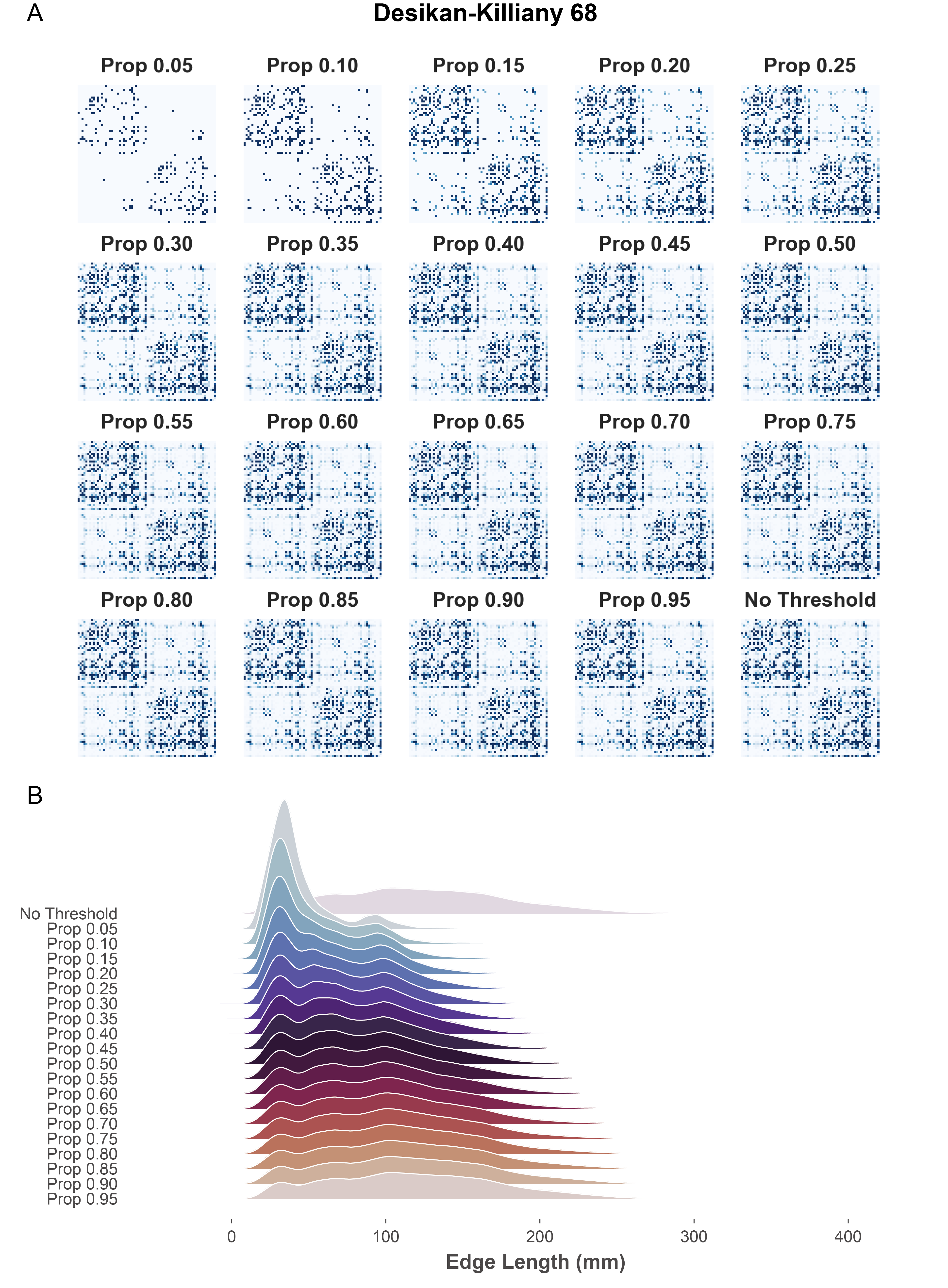
**Supplementary Fig. 3. Representative structural connectivity matrices and edge-length distributions based on Desikan-Killiany atlas.**

(A) Representative individual structural connectivity (SC) matrices based on the anatomically defined Desikan-Killiany atlas (68 nodes) across varying proportional thresholding strategies. Matrices are shown for proportional thresholds (Prop) ranging from 0.05 to 0.95, alongside the unthresholded baseline (No Threshold). (B) Density distributions of edge lengths (mm) for the connections retained under each thresholding strategy.

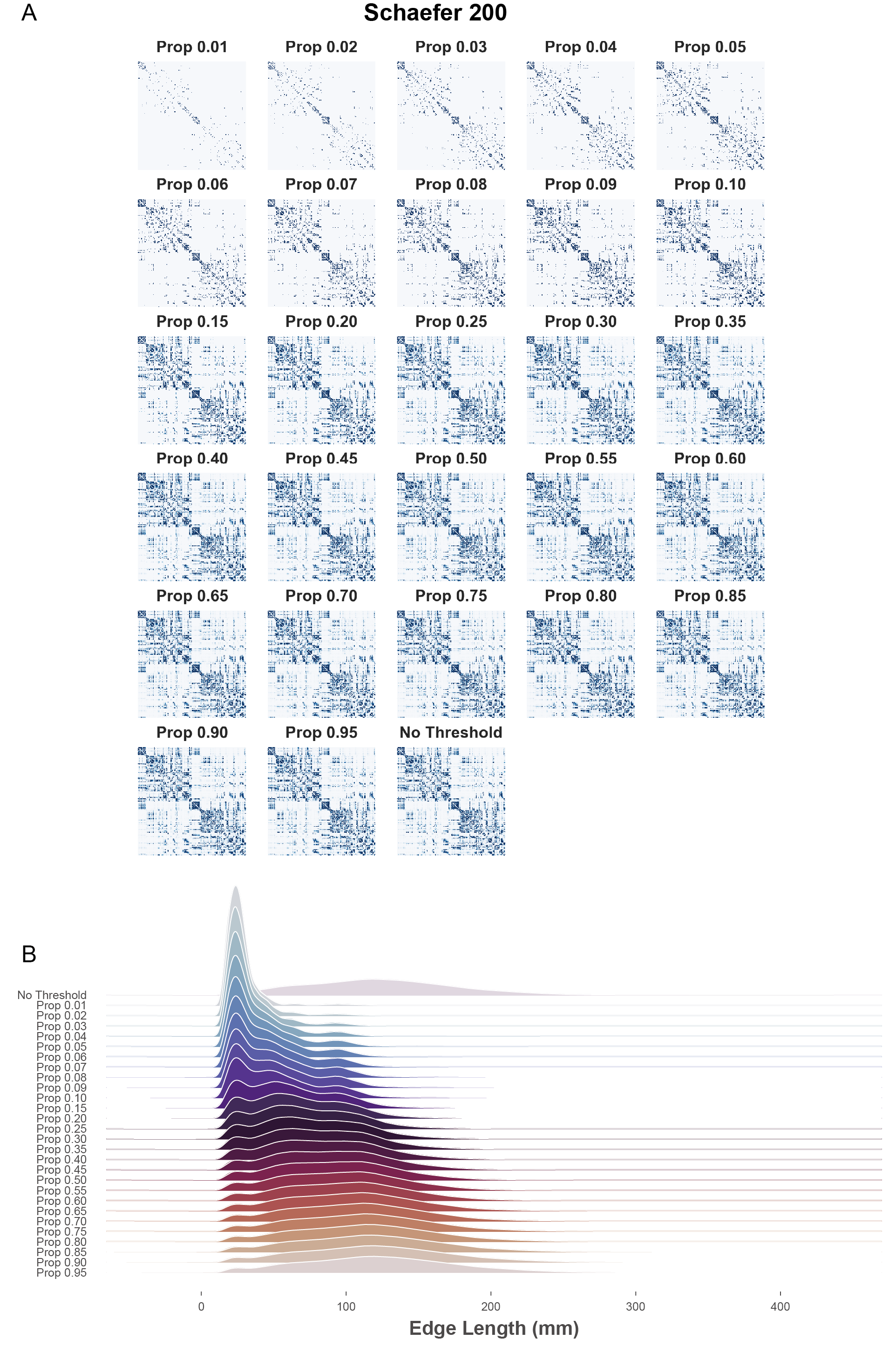

**Supplementary Fig. 4. Representative structural connectivity matrices and edge-length distributions based on Schaefer 200 atlas.**

(A) Representative individual structural connectivity (SC) matrices based on Schaefer 200 atlas (200 nodes) across varying proportional thresholding strategies. Matrices are shown for proportional thresholds (Prop) ranging from 0.01 to 0.95, alongside the unthresholded baseline (No Threshold). (B) Density distributions of edge lengths (mm) for the connections retained under each thresholding strategy.

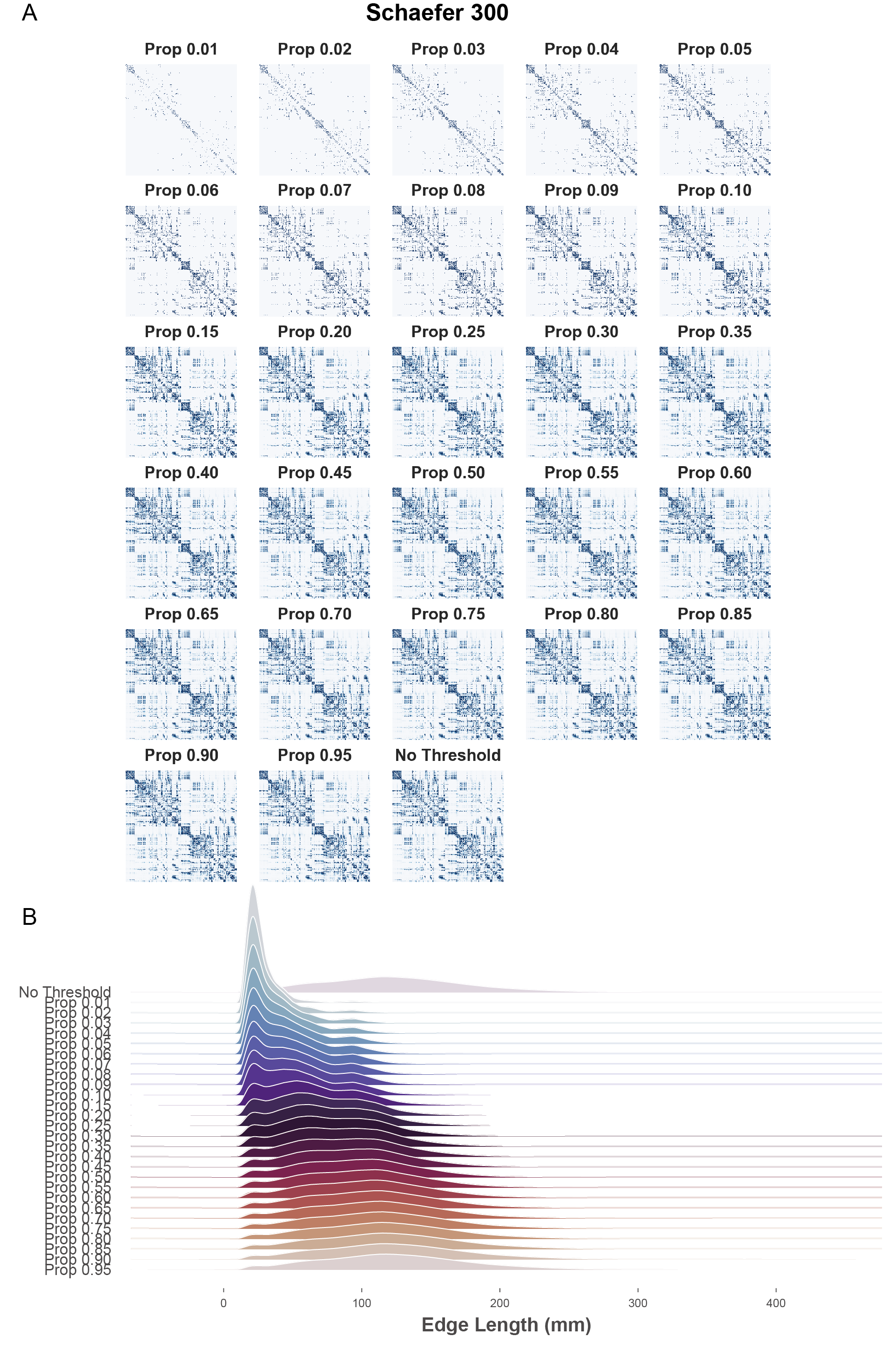

**Supplementary Fig. 5. Representative structural connectivity matrices and edge-length distributions based on Schaefer 300 atlas.**

(A) Representative individual structural connectivity (SC) matrices based on Schaefer 300 atlas (300 nodes) across varying proportional thresholding strategies. Matrices are shown for proportional thresholds (Prop) ranging from 0.01 to 0.95, alongside the unthresholded baseline (No Threshold). (B) Density distributions of edge lengths (mm) for the connections retained under each thresholding strategy.

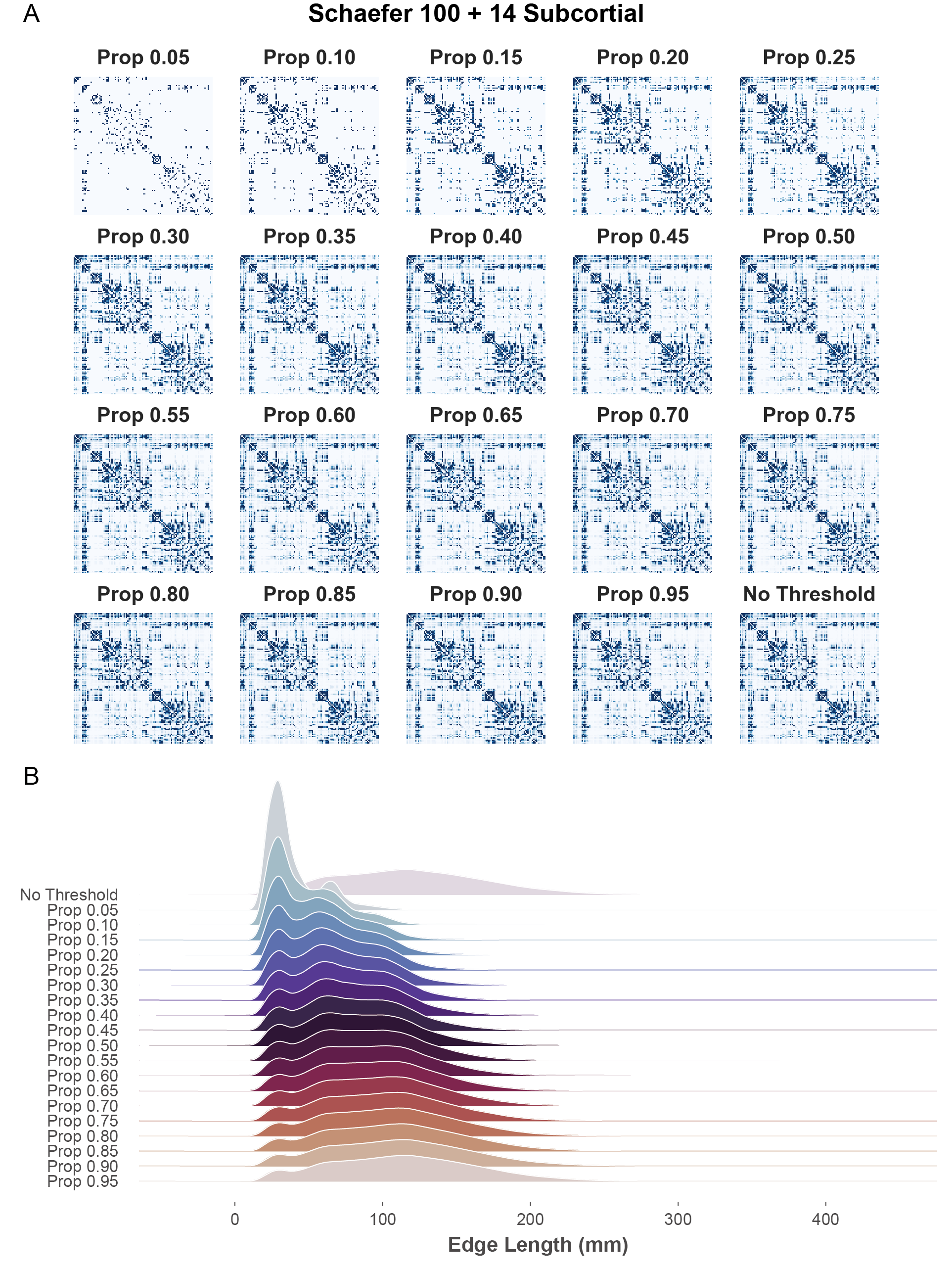

**Supplementary Fig. 6. Representative structural connectivity matrices and edge-length distributions based on Schaefer 100 plus 14 subcortical nodes.**

(A) Representative individual SC matrices based on the hybrid configuration (Schaefer 100 plus 14 subcortical nodes) across varying proportional thresholding strategies. Matrices are shown for proportional thresholds (Prop) ranging from 0.05 to 0.95, alongside the unthresholded baseline (No Threshold). (B) Density distributions of edge lengths (mm) for the connections retained under each respective thresholding strategy shown in (A).

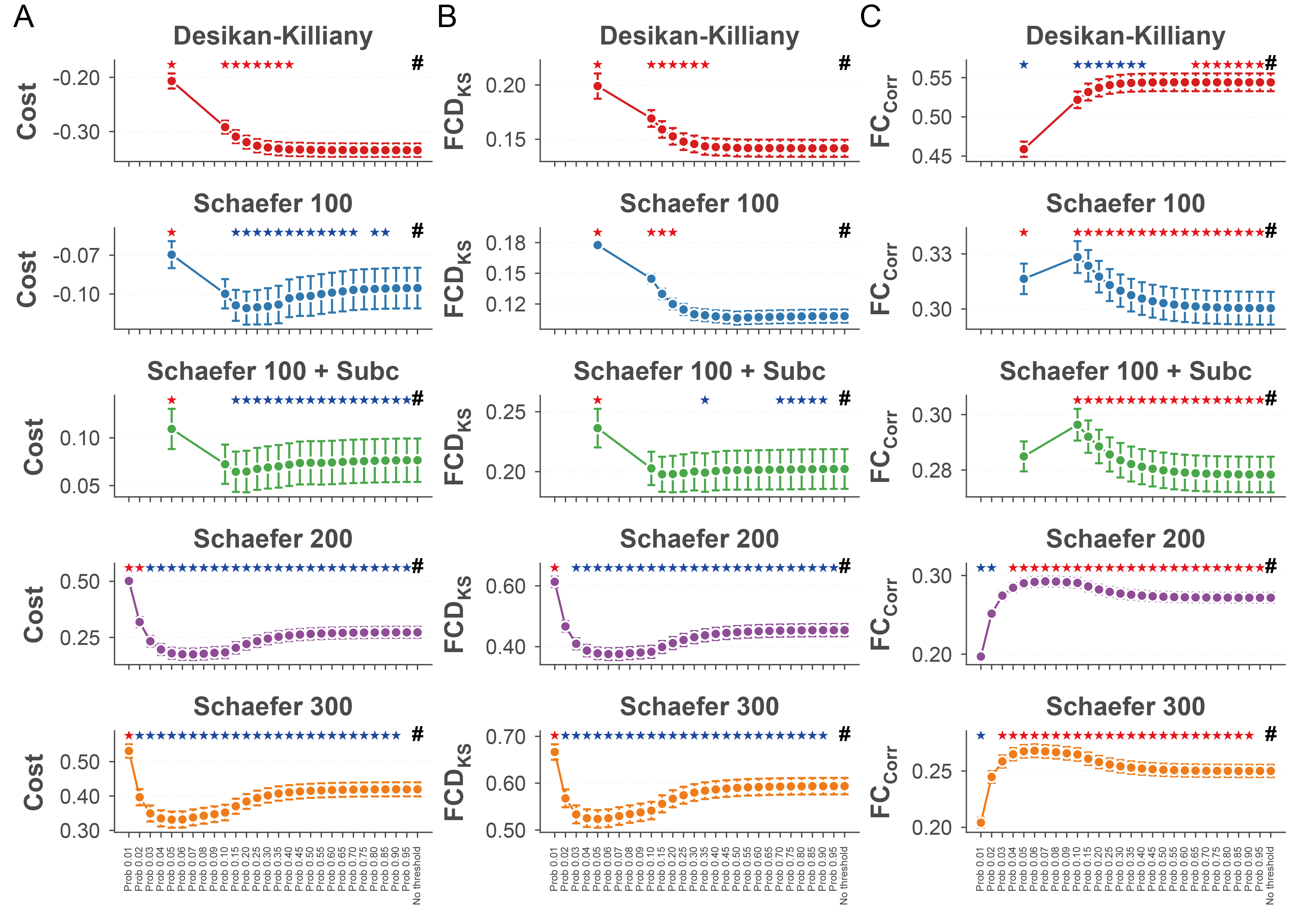

**Supplementary Fig. 7. Performance metrics of the DMF model across varying spatial scales and network densities.**

(A-C) Evaluation of model fitting across different parcellation granularities (Desikan-Killiany, Schaefer 100, Schaefer 100 + Subcortical, Schaefer 200, and Schaefer 300) and proportional thresholding densities. Performance is quantified by the non-normalized, raw values of the integrated $Cost$ metric (A), $FCD_{KS}$ (B), and $FC_{Corr}$ (C). The '#' symbol denotes the unthresholded baseline. Stars indicate statistically significant differences compared to the baseline; blue indicates a significant decrease, while red indicates a significant increase (P < 0.05, paired t-test). Data are presented as mean ± SEM.

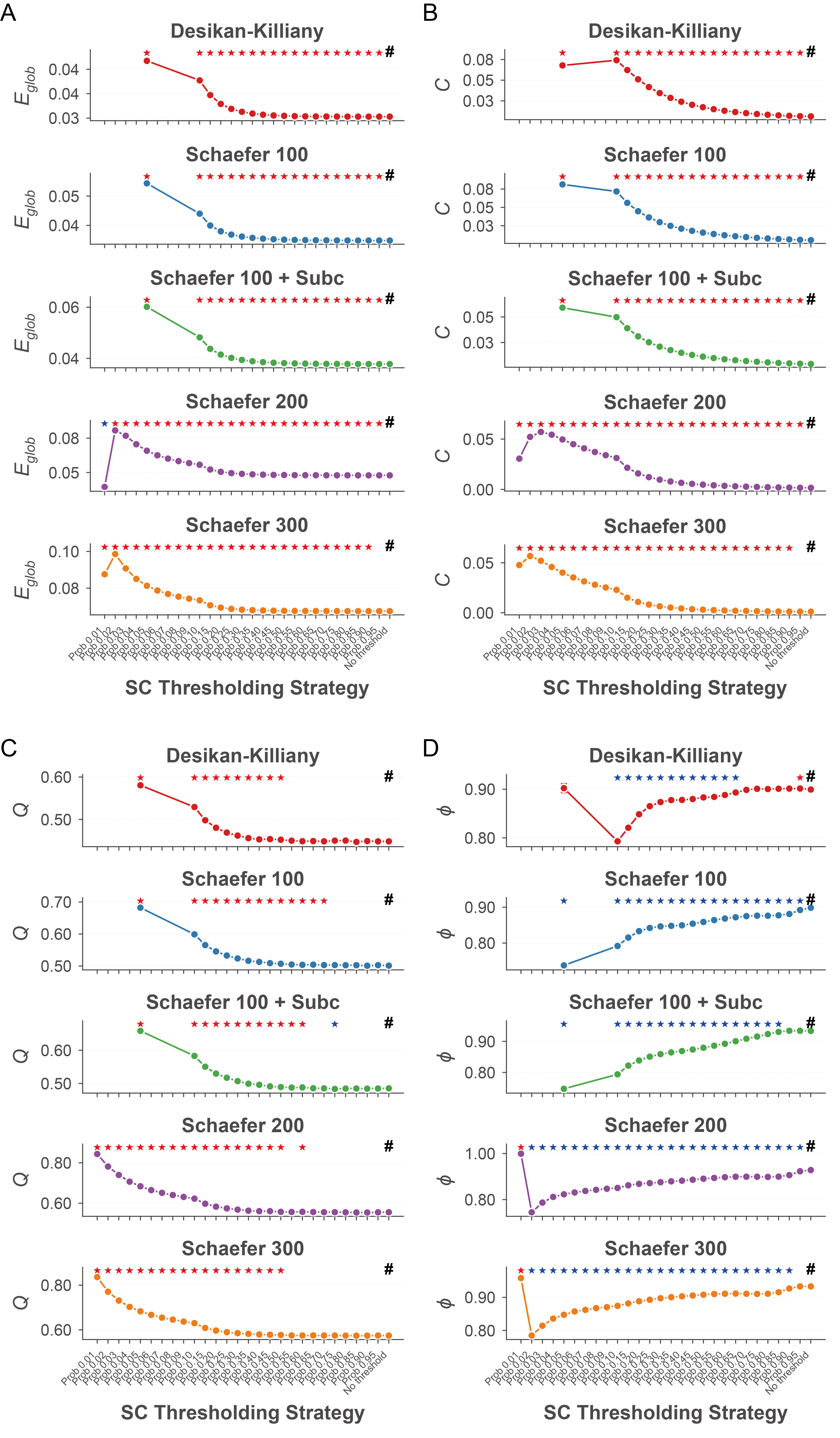
**Supplementary Fig. 8. Graph-theoretical topological metrics of structural connectivity networks across varying spatial scales and network densities.**

(A-D) Evaluation of graph-theoretical topological metrics for structural connectivity (SC) networks across different parcellation granularities (Desikan-Killiany, Schaefer 100, Schaefer 100 + Subcortical, Schaefer 200, and Schaefer 300) and proportional thresholding densities. The topological properties are quantified by the non-normalized, raw values of global efficiency $E_{glob}$ (A), clustering coefficient $C$ (B), modularity $Q$ (C), and small-worldness propensity $\phi$ (D). The '#' symbol denotes the unthresholded baseline. Stars indicate statistically significant differences compared to the baseline; blue indicates a significant decrease, while red indicates a significant increase (P < 0.05, paired t-test). Data are presented as mean $\pm$ SEM.
